# The chalk streams of southern England and northern France harbour substantial unique components of the overall genetic diversity of Atlantic salmon (*Salmo salar* L.)

**DOI:** 10.1101/2025.08.27.672665

**Authors:** R. Andrew King, Guillaume Evanno, Jamie R. Stevens

## Abstract

Populations of Atlantic salmon continue to suffer marked declines in abundance due to stressors acting in both their freshwater and marine habitats. It is therefore an imperative to identify populations in need of increased conservation intervention, with the aim of preserving as much as possible the genetic diversity present within the species. Previous microsatellite-based analyses have shown the chalk rivers of southern England and northern France to hold genetically distinct populations of salmon. However, these salmon populations have never been investigated in the same study. Using a suite of 93 single nucleotide polymorphism loci and samples from 42 British Isles and French rivers, we demonstrate the French and English chalk salmon to be closely related and distinct from salmon inhabiting non-chalk rivers. The identification of a small number of significant F_ST_ outliers suggests that this distinction is driven by local adaptation. We propose that the chalk and non-chalk salmon be designated as two distinct Evolutionarily Significant Units that each contain multiple Management Units. The chalk river salmon, especially those from southern England, are identified as making a significant contribution to the overall diversity of the species within the English Channel region. As a consequence, we propose that the salmon populations of the chalk streams may meet the criteria for recognition as a distinct subspecies of salmon, *Salmo salar calcariensis*. Taken together, the results presented here highlight the urgent need for enhanced conservation and protection for the Atlantic salmon populations inhabiting the chalk rivers of southern England and northern France.

## 1. Introduction

Freshwater biodiversity is in a state of global crisis (Dudgeon, 2019). Increasingly it is becoming clear that anthropogenic activities are having adverse effects on a wide range of ecological and evolutionary processes, leading to drastic and accelerating declines in size of wildlife populations (WWF, 2024). The consequences of these activities are being felt in all terrestrial and aquatic habitats but especially so in freshwater habitats (Dudgeon, 2019; WWF, 2024) through the multiple, co-occurring stressors of habitat degradation and loss, both point and diffuse source pollution, exploitation of water resources, i.e. over abstraction, invasive species and climate change (Dudgeon, 2019; Haase et al., 2025). The negative effects of stressors on species and ecosystems can lead to losses of genetic diversity, changes in species ranges and species extinctions (Comte et al., 2013; Exposito-Alonso et al., 2022; Shaw et al., 2025; Urban, 2024). Globally, populations of migratory fish have shown particularly striking declines in abundance with reductions in excess of 80% over the last 50 years, with the most marked declines occurring since the mid-1990s (Deinet et al., 2024). Within Europe, species abundances have declined by 75% since the mid-1990s. Atlantic salmon (*Salmo salar*) is an iconic anadromous fish species. After spending one to several years in freshwater, salmon smoltify and migrate to sea to feed and grow, usually for one to three years, before returning to their natal river to spawn (Crisp, 2000; Klemetsen et al., 2003). The species has suffered marked range-wide declines due to the combined effects of stressors in both its freshwater natal and spawning habitat and its marine feeding habitats (Dadswell et al., 2022; Gillson et al., 2022; Nunn et al., 2023). Recently, the IUCN have updated the conservation status of Atlantic salmon, downgrading the species globally to near threatened. Assessments were also published for different genetic groups within the species, with the salmon populations of Britain classified as endangered, the English chalk stream populations as vulnerable, the French Allier sub-population as endangered and the France/Spain grouping as of least concern (Darwell, 2023a, 2023b, 2023c, 2023d; Evanno et al. 2023).

In a natural world increasingly affected by the detrimental actions of humans, it is important to identify populations in need of increased conservation management and to safeguard as many populations as possible to meet recent United Nations targets of protecting at least 90% of diversity present within a species (Exposito-Alonso et al., 2022). Genetic and genomic data have been used extensively to help inform conservation management decision-making processes (Forrester & Lama, 2023). For some species, the conservation of genetically distinct populations, which can represent unique evolutionary lineages, has been prioritised (Chhina et al., 2024; von Takach et al., 2024).

Chalk streams are a globally rare habitat found predominately in southern and eastern England but also in northern France and Denmark (CABA, 2021; WWF-UK, 2015). Such watercourses are dominated by groundwater flowing from Cretaceous-age chalk aquifers (Berrie, 1992; Sear et al., 1999). The chalk has a strong influence on the flow regime, temperature and chemical properties of the river water resulting in chalk streams supporting characteristically distinctive plant and animal assemblages (Berrie, 1992; Sear et al., 1999).

Several studies have shown that the salmon populations inhabiting the chalk streams of the Hampshire Basin in southern England are one of the most genetically unique groups of rivers in the north east Atlantic distribution of Atlantic salmon (Finnegan et al., 2013; Gilbey et al., 2018; Griffiths et al., 2010; Ikediashi et al., 2018). Likewise, in northern France, the salmon populations from the rivers of Upper Normandy, which also flow over chalk substrate, are among the most distinct within the French distribution of the species (Perrier et al., 2011). These studies, based on analysis of microsatellite loci, have generally shown lower levels of genetic diversity in chalk rivers compared to neighbouring rivers flowing over non-chalk substrates (Finnegan et al., 2013; Ikediashi et al., 2018; Perrier et al., 2011). However, the Hampshire Basin and Upper Normandy salmon populations have never been investigated in the same study. Moreover, being outside the main range of Atlantic salmon in western Europe (Scotland, Ireland and Norway) coverage of English Channel populations has been sparse in recent range-wide single nucleotide polymorphism (SNP) based studies (Bradbury et al., 2021; Jeffery et al., 2018; O’Sullivan et al., 2022).

Here, we explore patterns of genetic diversity in Atlantic salmon from 42 rivers from the southern British Isles and northern France using a recently developed panel of SNP markers (King & Stevens, 2021), with an emphasis on populations in the chalk rivers of the Hampshire Basin and Upper Normandy. Our objectives were to 1) assess patterns of genetic diversity and structuring of salmon within these rivers, 2) determine the relationship between salmon inhabiting the chalk rivers of southern England and northern France, and populations in non-chalk rivers, and 3) identify salmon populations in need of targeted conservation management.

## 2. Materials and Methods

### 2.1 Sample collection

Individual Atlantic salmon were sampled from 42 English, Irish and French rivers (Figure 1, Supplementary Table 1). The majority of fish were caught as fry or parr during routine management surveys by the Environment Agency, Natural Resources Wales, Inland Fisheries Ireland and the Game and Wildlife Conservation Trust. Collections typically consisted of samples collected during electrofishing surveys from multiple sites across each river catchment. Fish were anaesthetised using 2-phenoxyethanol or MS-222 prior to removal of tissue samples (either fin clips or scales). Fin clips were transferred immediately into tubes containing absolute ethanol. Scale samples were obtained from trap- or rod-caught adults from the Tamar, Wye and northern French rivers. Genomic DNA was extracted from fin clips following the method of Truett *et al*. (2000) and scales using Qiagen DNeasy Blood and Tissue kits, following the manufacturer’s instructions.

**Figure 1.**
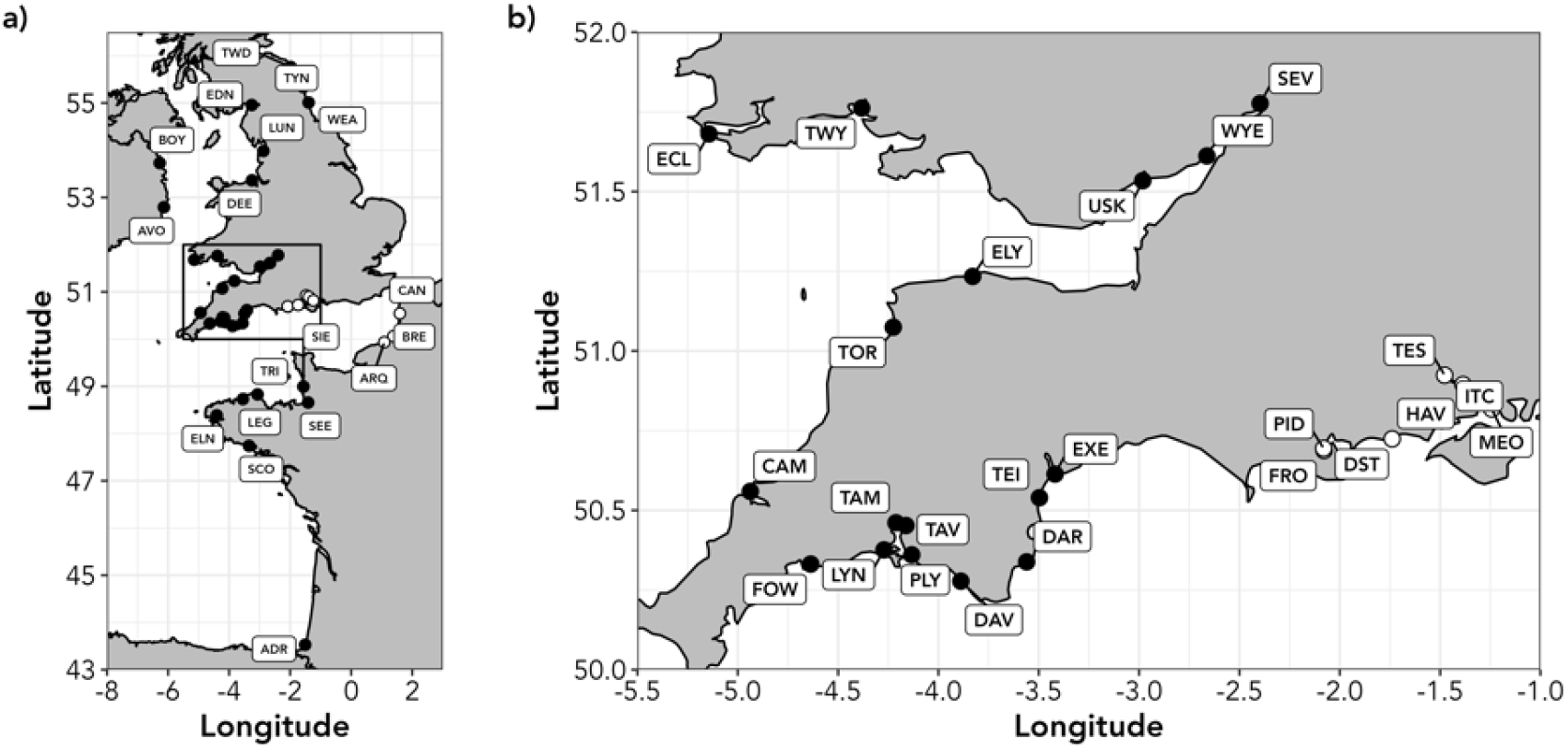
Map showing the sampled Atlantic salmon rivers in Britain, Ireland and France: a) location of the 42 sampled rivers, b) detail of rivers in southern England and Wales. For clarity, only the location of the river mouth is shown. White and black symbols represent chalk and non-chalk rivers, respectively. River codes are as given in Supplementary Table 1.

### 2.2 SNP Genotyping

Each sample was screened for variation at 94 biallelic single nucleotide polymorphism (SNP) markers recently developed for Atlantic salmon (King & Stevens, 2021). SNP genotyping was undertaken on a Standard Biotools EP1 Genotyping System using 96.96 Dynamic Genotyping Arrays and scored using the Standard Biotools SNP Genotyping analysis software. Genotype plots were manually inspected to ensure quality of individual genotyping and clustering. Each genotyping run included two positive (individuals of known genotype) and two negative controls.

### 2.3 Data quality control

Salmonid populations can often contain numbers of closely related individuals (Hansen et al., 1997), the presence of which can potentially lead to bias in population genetic parameter estimation (Goldberg & Waits, 2010) and false inference of population structure (Rodríguez-Ramilo & Wang, 2012). For sibship reconstruction, we used COLONY v 2.0.5.9 (Jones & Wang, 2010), which implements a maximum-likelihood method to assign sibship to individuals using their multilocus genotype. Run conditions were: high precision, medium length run, assuming both male and female polygamy without inbreeding, and a conservative 0.5% error for both scoring error rate and allelic dropout rate. Fish were considered members of a full-sib family if the probability of exclusion was > 0.9. We used the Waples & Anderson (2017) Yank-2 method to trim full sibs from the data set – both members of families with two individuals were retained but if a family had three or more individuals, all but two random members were removed.

Genepop v 3.4 (Raymond & Rousset, 1995) was used to test for linkage disequilibrium (LD) between all pairs of loci within each population. Significance was estimated using a Markov-chain method (1000 de-memorisations, 100 batches and 1000 iterations). Hochberg’s (1988) correction was used to adjust significance levels for multiple comparisons (https://www.multipletesting.com, Menyhart et al. (2021)). Deviation from Hardy-Weinberg Equilibrium (HWE), based on Nei’s (1987) heterozygosity-based G_IS_ estimator, for each locus and population was tested using GenoDive v3.03 (Meirmans, 2020). Significance was based on 999 permutations.

### 2.4 Genetic diversity

Basic measures of genetic diversity (H_O_ - observed heterozygosity and H_S_ - unbiased expected heterozygosity) and Weir & Cockerham’s (1984) estimator of F_ST_ were calculated using GenoDive v3.03 (Meirmans, 2020). Significance of F_ST_ values was assessed using 999 bootstrap replicates. The percentage of polymorphic loci in each collection after subsampling to the smallest number of genotyped fish was calculated using the R scripts of Anderson et al. (2025).

### 2.5 Selection tests

#### 2.5.1. F_ST_-based outlier tests

Two F_ST_-based tests were performed to identify loci under selection. Bayescan 2.01 (Foll & Gaggiotti, 2008) implements the Bayesian approach of Beaumont & Balding (2004). The program was run under default settings (20 pilot runs of 5000 iterations each followed by an additional burn-in of 50 000 iterations and then 5000 samplings with a thinning interval of 10). We complemented the Bayescan results with simulations using the fdist model as implemented in Arlequin 3.5 (Excoffier & Lischer, 2010) using default parameters (10 000 simulations with a null model with 10 groups, each containing 100 demes). Both Bayescan and fdist analyses were run on three separate datasets – full (42 rivers, 93 SNPs), non-chalk (32 rivers, 89 SNPs) and chalk (10 rivers, 88 SNPs) datasets. Additionally, hierarchical genetic structuring can lead to an excess of false positives if not accounted for in outlier tests (Excoffier, Hofer, & Foll, 2009). We therefore repeated the fdist analysis on the full dataset using the hierarchical F_ST_ model based on two data partitions: chalk v non-chalk, and UK chalk v FR chalk v UK non-chalk v FR non-chalk. Loci were considered under selection if they were outside of the 99% confidence level of the simulated neutral distribution.

#### 2.5.2 Redundancy Analysis

To investigate associations between genetic data and environmental variables we performed a Redundancy Analysis (RDA), implemented in the *vegan* v2.5-2 R package (Oksanen et al., 2019). Genetic data was the minor allele frequency (MAF) at each SNP locus. Environmental data for each river, in the form of 19 bioclim variables at 1km spatial resolution, were extracted from WorldClim databases using the *raster* v3.0-7 R package (Fick & Hijmans, 2017; Hijmans, 2019). For rivers where fish were sampled from multiple sites, we used the data for furthest downstream site. For the northern French rivers, where samples were collected from rod-caught adults from across each catchment, we choose a point approx. 10km upstream of the river mouth as the environmental site. To account for significant correlations between the bioclim variables we initially ran the RDA with all 19 variables and progressively removed the variable with the highest variance inflation factor (vif) until all remaining vif values were less than 10. Analysis of Variance (ANOVA), based on 999 permutations, was used to test the significance of the full RDA model, each RDA axis and each bioclim variable. Candidate SNP loci were identified in the tail of the allele loading distribution for each axis, using a 3-standard deviation (SD) from the mean loading cut-off (corresponds to a two-tailed *p*-value = 0.0027 - Forester *et al*. 2018). A second RDA was run conditioned on geographical distance from the northern-most river (the Tweed). Least-cost marine distances between the mouth of each river and the Tweed were calculated using the marmap R package (Pante & Simon-Bouhet, 2013).

For SNP loci identified as outliers in F_ST_-based tests and showing significant associations with environmental data we identified the genome location of each locus. The original RADtag sequences (King & Stevens, 2021) were aligned to the Atlantic salmon genome (Ssal_v3.1) using the BLAST option in SalmoBase (Samy et al., 2017). We noted linkage group, location and whether the SNP was present in a gene or non-coding region.

### 2.6 Genetic structure

To investigate genetic structuring, we performed three analyses. Firstly, using POPULATIONS v1.2.32 (Langella, 1999), a neighbour-joining dendrogram based on Cavalli-Sforza and Edwards (1967) chord distance (D_CE_) was constructed and visualised using MEGA v6 (Tamura, Stecher, Peterson, Filipski, & Kumar, 2013). Secondly, data were analysed using the Bayesian-based Markov Chain Monte Carlo (MCMC) model-based clustering method implemented in STRUCTURE v 2.3.4 (Pritchard et al., 2000), which jointly defines *K*, the number of possible partitions of the data set and the proportion of an individual’s genome (q) derived from each of the *K* partitions. STRUCTURE was run for 250 000 iterations following a burn-in of 100 000 iterations with the number of inferred populations (*K*) ranging from one to ten. Ten independent runs at each *K* were performed using the admixture model with correlated allele frequencies and not using the river of origin information as a prior. The most likely number of clusters was determined using the Δ*K* method of Evanno *et al*. (2005). To identify finer-levels of structure, hierarchical analyses were performed based on the Δ*K* results for the full data set. Consensus data were visualised using POPHELPER v1.0.6 (Francis, 2017). Finally, Discriminant Analysis of Principal Components (DAPC; Jombart et al. (2010)) analyses were undertaken using the *adegenet* (Jombart, 2008) package for R (R Core Team, 2018). The optim.a.score() function was used to select the optimum number of principal components to be retained in the analysis. As with the STRUCTURE analysis, hierarchical analyses were performed based on the results for the full dataset. For hierarchical analyses, loci found to be near monomorphic in either chalk or non-chalk rivers were removed (five and four loci, respectively) from the datasets (Supplementary Table 2).

### 2.7 Identification of Conservation Units and Conservation priority

A key aim of conservation genetics is to highlight populations within a species that may require special conservation measures. We used multiple approaches to identify groups of rivers in need of urgent conservation intervention. Firstly, using the three-step approach of Funk *et al*. (2012), we delineated Conservation Units (CUs) for the sampled rivers. Step 1 uses the information from all loci to identify Evolutionarily Significant Units (ESUs). Then, using data from neutral loci, demographically independent Management Units (MUs) were defined. Finally, adaptively differentiated units (AUs) were recognised using outlier loci. To determine the population groupings, we constructed population-based neighbour-joining dendrograms, as described above, based on 93 loci for the full, 81 loci for the neutral and 12 loci for the outlier datasets, respectively. The outlier dataset contained genotypes for the 12 loci found to be under selection in the Bayescan, fdist and fdist hierarchical analyses.

Finally, we estimated the expected contribution of each river to two diversity measures - gene and allelic diversity using the approaches of Caballero & Rodríguez-Ramilo (2010) and Petit *et al*. (1998) as implemented in METAPOP2 (López-Cortegano, Pérez-Figueroa, & Caballero, 2019). The contribution of a river to within subpopulation (*H*_S_, *A*_S_), between subpopulation (*D*_G_, *D*_A_) and total (*H*_T_, *A*_T_) gene and allelic diversity, respectively, was estimated by serial removal of each river from the dataset and calculating the change in each diversity metric. Rivers can have either a negative or positive contribution to each diversity measure with a positive contribution signifying a loss of diversity after removal of a river and an increase in diversity indicating a negative contribution. We also calculated each diversity measure for the four main population groupings identified in our genetic structure analyses.

## 3. Results

### 3.1 Data quality

A total of 1640 individuals were successfully genotyped at a minimum of 89 out of the 94 SNP assays. Comparison of genotypes from repeated samples gave an error rate of 0.0011% (ten mismatches from 9024 allele calls). All loci were polymorphic in at least ten of the sampled rivers, with levels of polymorphism ranging from 81.7% to 93.4% (Supplementary Table 1). COLONY analysis found a total of 100 full sib families (range 0-9 families per sample and 2-19 members per family). A total of 75 full sibs were trimmed, leaving a final data set of 1565 salmon on which all subsequent analyses were performed.

Tests found a total of 34 significant cases of linkage between pairs of markers, with the majority in chalk rivers. Twelve and six of these cases involved marker pairs Ssa_25077 - Ssa_1354 and Ssa_25077 - Ssa_41749, respectively. Consequently, the data for Ssa_25077 were removed, leaving a final data set of 93 SNP markers.

There were 142 significant deviations from HWE (out of a total of 3906 river/locus combinations). None of these significant results were consistent across loci or rivers. Seven rivers showed significant deviations from HWE (Supplementary Table 1), with three rivers (Meon, Sienne and Scorff) showing a significant heterozygote deficiency and four (Wear, Tamar, Dart and Stour) showing a significant excess of heterozygotes. Overall, all rivers and remaining loci were retained for further analyses.

### 3.2 Genetic diversity

Basic measures indicated marked differences in genetic diversity between chalk and non-chalk salmon rivers (Supplementary Table 1). Both observed and expected heterozygosity were generally higher in the western non-chalk and French chalk rivers and lowest in the Hampshire Basin chalk stream populations (Supplementary Table 1). The lowest H_S_ values were found in salmon in the East Lyn and Meon (Supplementary Table 1) – two rivers with very small salmon populations. After accounting for sample size, the Dorset Stour had the lowest percentage of polymorphic loci while the highest values were found in the Upper Normandy rivers (Supplementary Table 1).

## 3.4 Loci under selection

### 3.4.1 FST-based outlier tests

Selection tests found a small number of loci under divergent selection in the three data sets. For the full data set, Bayescan and fdist found two and 11 loci, respectively, under divergent selection, with only a single locus (Ssa_69865) common to both analyses (Figure 2). Eleven of the outlier loci showed clear frequency differences between chalk and non-chalk rivers (Supplementary Figure 2), with the remaining locus (Ssa_six6) demonstrating high MAF in the Bristol Channel rivers (Usk, Wye & Severn). For the non-chalk rivers, Bayescan and fdist found one and three loci, respectively, under divergent selection, with only a single locus (Ssa_six6) common to both analyses (Supplementary Figure 1). For the chalk-only data set outlier loci were only found in the fdist analysis with five loci identified as outliers (Supplementary Figure 1). Considering multiple groups of rivers, the hierarchical fdist F_ST_ model found two SNPs, including Ssa_69865, under divergent selection in the analysis considering four groups of rivers (Supplementary Figure 3). Both SNPs showed striking allele frequency differences between the chalk and non-chalk river populations, being polymorphic in chalk rivers but near monomorphic in non-chalk salmon populations (Supplementary Figure 2).

**Figure 2.**
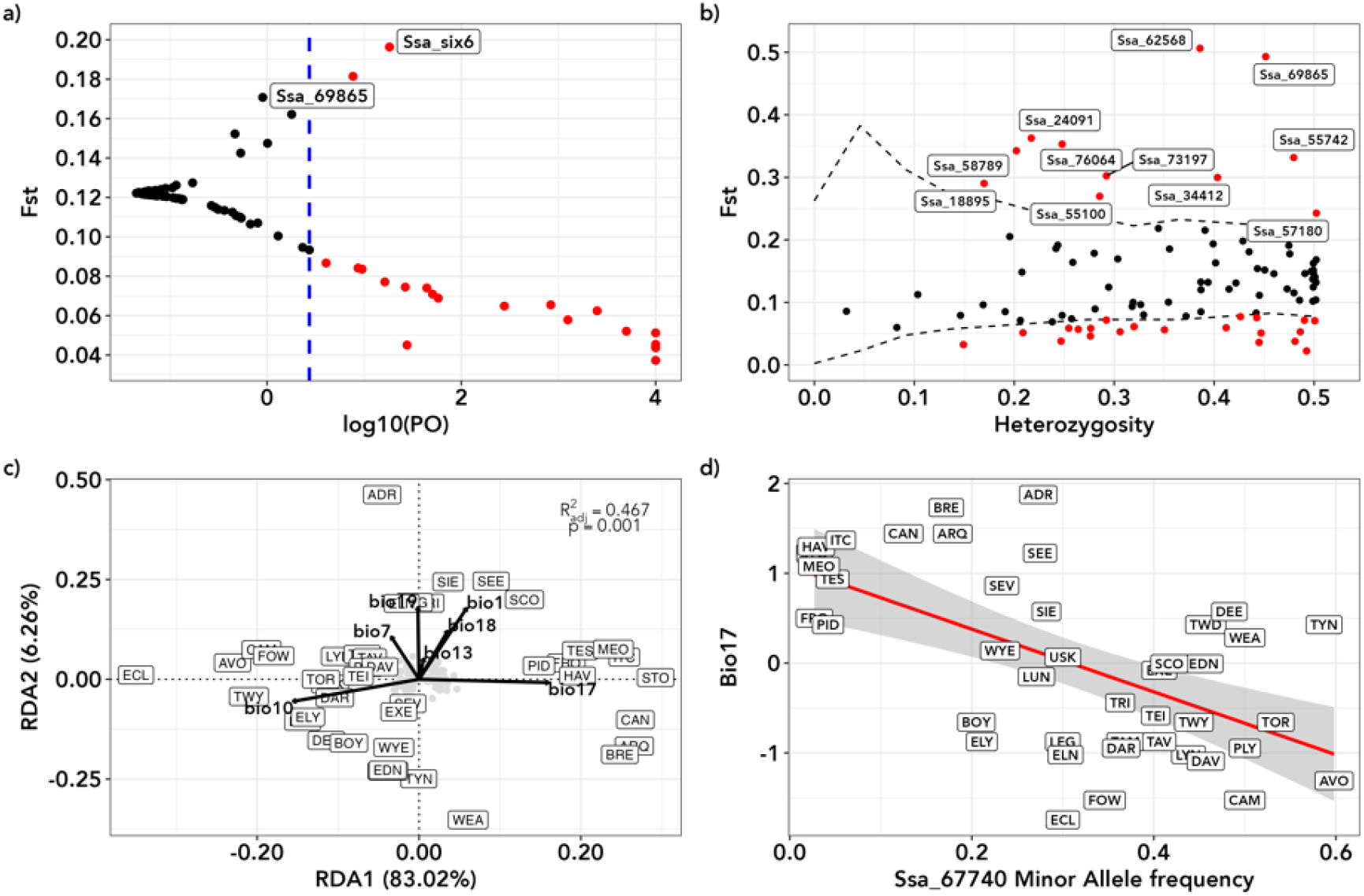
Results of selection and environmental association tests a) Bayescan. The dashed blue line represents the log_10_ PO value above which loci are considered significant ouliers. b) fdist results. Dashed lines are the upper and lower 99% confidence level of the simulated neutral distribution. Loci under either divergent or balancing selection are shown in red with loci under divergent selection labelled with locus name. c) Plot of unconditioned redundancy analysis axes 1 v 2. Black vectors represent loadings for the retained bioclim variables. d) Plot of minor allele frequency for locus Ssa_67740 versus standardised bio17 (precipitation of the driest quarter) values. The red line represents the linear regression, and the grey shaded area is the 95% confidence interval. For c) and d) river codes are as given in Supplementary Table 1.

**Figure 3.**
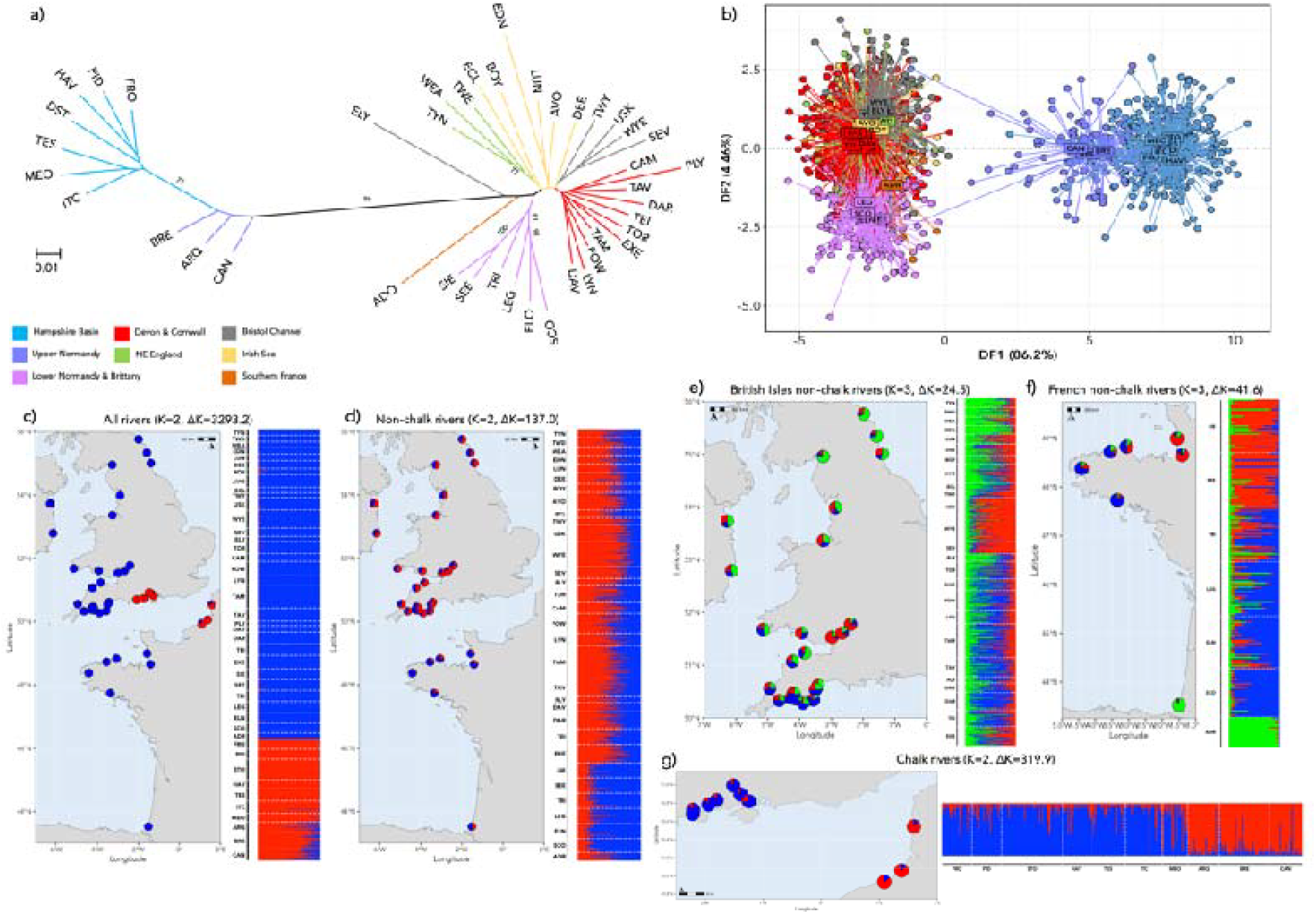
Results of the genetic structuring analyses for Atlantic salmon sampled from 42 British, Irish and French rivers. a) unrooted neighbour-joining dendrogram, based on Cavalli-Sforza and Edwards’ chord distance; branches are colour coded by geographical location. b) Discriminant Analysis of Principal Components analysis for 1565 Atlantic salmon showing discriminant function 1 v 2; each dot represents a sampled individual fish and are coloured as in a). c-g) Results for hierarchical STRUCTURE analyses; results of each analysis are shown as bar plots with horizontal columns representing the assignment probabilities of individuals to each of the K inferred clusters. Maps show the location of each sampled river mouth with pie charts giving the river-level assignment to each genetic cluster. Plots of ΔK values for STRUCTURE analyses are given in Supplementary Figure 6. River codes are as given in Supplementary Table 1.

The 12 F_ST_ outlier loci were found on ten linkage groups with eight of the loci being located in introns of known genes (Supplementary Table 3). Three loci were present in non-coding regions and a single locus was found in a putative pseudogene.

### 3.4.2 Redundancy Analysis

For the final model seven bioclim variables with vif values under ten were retained; temperature variables bio1, bio7 and bio10 and precipitation variables bio13, bio17, bio18 and bio19. Both unconditioned and geographic distance conditioned analyses gave very similar results. In both cases the full model was highly significant (unconditioned - F=5.20, p=0.001, adjusted r^2^=0.467; conditioned - F=5.43, p=0.001, adjusted r^2^=0.409). Axis 1 separated the chalk rivers from non-chalk rivers with this distinction being driven by mean temperature of the warmest quarter (bio10) and precipitation of the driest quarter (bio17). Axis 2 highlighted the Lower Normandy, Brittany and southern French rivers with this distinction being driven mainly by annual mean temperature (bio1) and precipitation of the coldest quarter (bio19) (Figure 2, Supplementary Figure 4).

**Figure 4.**
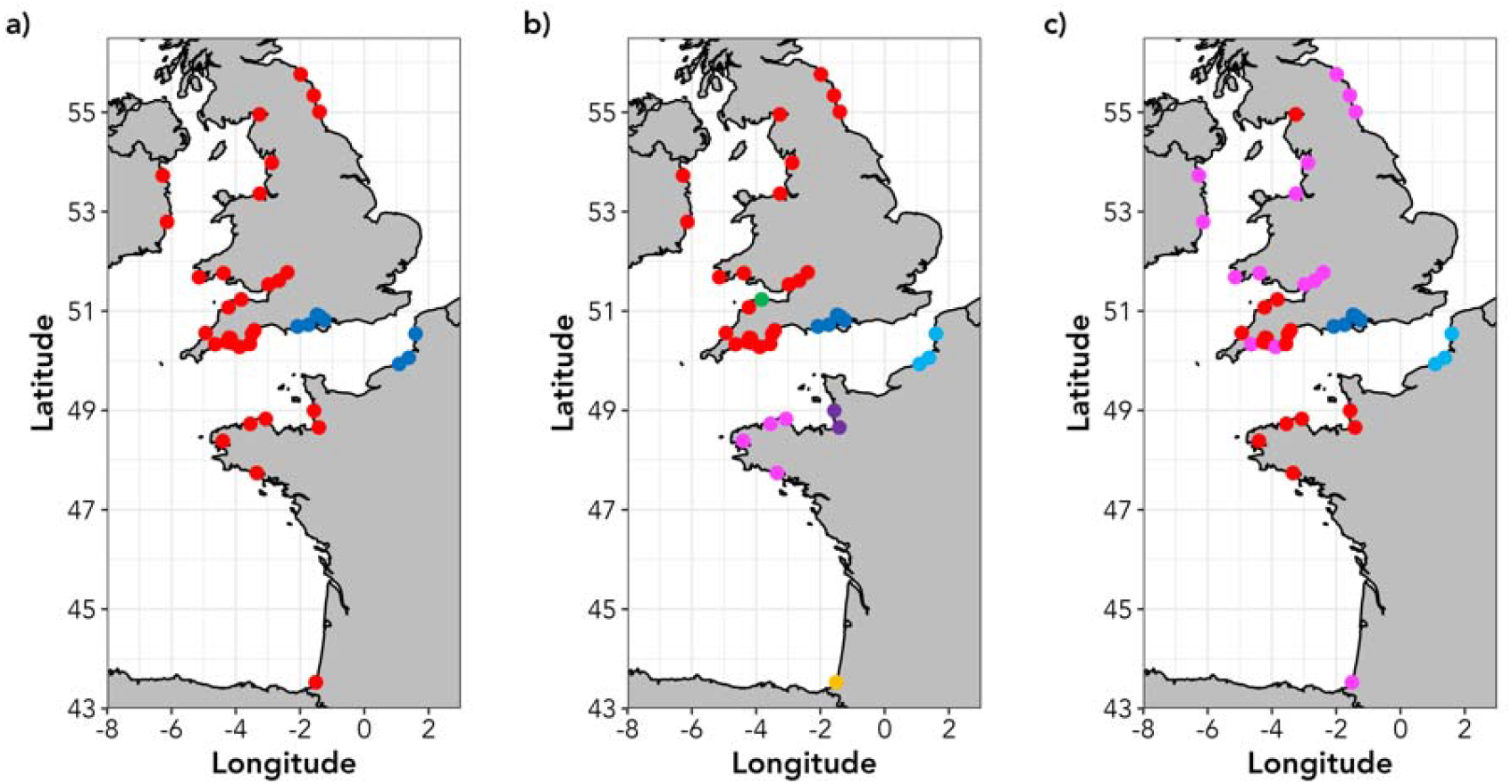
Maps showing designation of conservation units in 42 Atlantic salmon rivers. a) distribution of Evolutionarily Significant Units, based on all loci datasets b) Management Units, based on neutral loci datasets. c) distribution of Adaptive Units based on the outlier loci dataset.

For the unconditioned analysis a single locus, Ssa_67740, was found to be associated with bio17 (Figure 2). The conditioned analysis also identified the Ssa_67740–bio17 association along with two additional loci – Ssa_30724 and Ssa_87179 being associated with bio17 and bio10, respectively (Supplementary Figure 4). Of the three SNPs associated with environmental variables only one (Ssa_67740) was located in a known gene, Ssa_87179 was located in an uncharacterised gene and Ssa_30724 was found in a non-coding region (Supplementary Table 3).

## 3.3 Genetic structuring

Population pairwise F_ST_ (Supplementary Table 4, Supplementary Figure 5) values ranged from −0.003 (Sienne v Sée) to 0.326 (Eden v Avon). All but five pairwise values where highly significant (Supplementary Table 4). Within the non-chalk river populations, average pairwise F_ST_ was 0.044 (range −0.003 – 0.141) despite very large geographic distances between some rivers (*i.e*., Tweed – Adour, least-cost geographic distance = 1761km; F_ST_ = 0.08). Within the chalk group average pairwise F_ST_ was 0.033 (range 0.002 – 0.067). Between non-chalk and chalk rivers, pairwise differentiation values were all greater than 0.145 (average = 0.254, maximum = 0.326).

**Figure 5.**
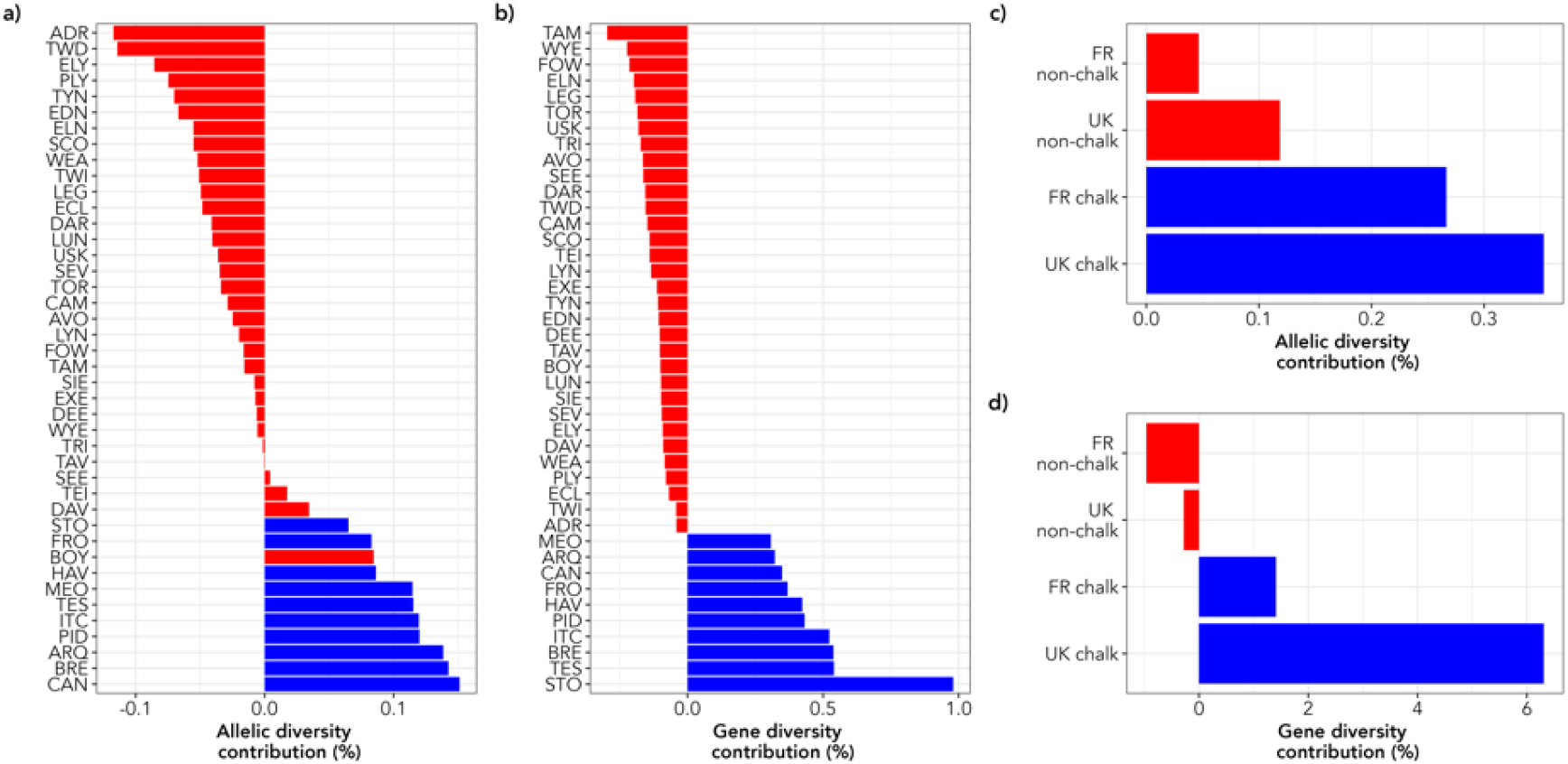
Conservation priorities for 42 Atlantic salmon rivers and four major regional groups of rivers. a – d) Contribution of 42 rivers (a & b) and four regional groups of rivers (c & d) to total allelic (A_T_) and gene (H_T_) diversity. Rivers/regions are sorted from lowest to highest contribution with non-chalk rivers shown in red and chalk rivers in blue.

The results of the neighbour-joining, STRUCTURE and DAPC analyses were in broad agreement. All three analyses confirmed the strong split between non-chalk and chalk river populations seen with pairwise F_ST_ values (Figure 3). For the STRUCTURE analysis, the Δ*K* method identified *K*=2 as the most likely partition of the full data set (Δ*K*=3293.2 – Supplementary Figure 6), differentiating the salmon from the chalk rivers of the Hampshire Basin and Upper Normandy rivers from all others. There was significant admixture between the non-chalk and chalk groups within the Arques, Bresle and Canche rivers, with an average ancestry contribution of 0.178 from the non-chalk group to Upper Normandy individuals. Hierarchical analyses identified further substructure within the two main groups of rivers. Within the non-chalk rivers there were multiple, distinct groups of populations. For instance, within the French non-chalk rivers, three groups were identified – Lower Normandy, Brittany and southern France (Figure 3). Within the British and Irish non-chalk rivers, the inner Bristol Channel rivers (Twyi, Severn, Wye and Usk) were distinct from other geographically proximate southwest Britain rivers (i.e. Camel, Torridge and Eastern Cleddau). The chalk rivers of the Hampshire Basin and Upper Normandy were distinct groups with little evidence of structuring within each region (Figure 3).

DAPC analysis showed the same strong split as the STRUCTURE analysis (Figure 3, Supplementary Figure 7). Within each of the two main clusters of individuals, there is separation between the British and French rivers (Supplementary Figure 6). This analysis also highlighted that two of the adult salmon sampled from Upper Normandy rivers (one each from the Arques and Canche) seem to be strays from the western non-chalk cluster (Supplementary Figure 8). The stray into the Arques (ARQ11) appears to have originated from either a Lower Normandy or a Breton river, while the Canche fish (CAN30) is likely a stray from the one of the southwest English rivers. The chalk-only DAPC shows a weak split between the eastern (Test, Itchen and Meon) and western (Frome, Piddle, Avon, Stour) Hampshire Basin rivers (Supplementary Figure 7) – a distinction that is not apparent in the STRUCTURE results.

## 3.6 Identification of Conservation Units

The population-based NJ dendrograms identified multiple groups of rivers that could be designated as distinct conservation units (Figure 4, Supplementary Figure 9). Conservatively, based on the data for all 93 loci, we identified two main groupings of rivers that could be classified as ESUs (Figure 4a, Supplementary Figure 9a). The neutral-only loci datasets identified seven groupings that could be designated as MUs, namely Upper Normandy, Lower Normandy, Brittany, Hampshire Basin, and western and northern British Isles, with two distinct rivers (ELY and ADR) potentially being single river MUs (Figure 4b, Supplementary Figure 9b). We distinguish the Brittany and Lower Normandy MUs on the basis of previously published microsatellite data (Perrier et al., 2011). The outlier loci again identified four main groupings. The Hampshire Basin and Upper Normandy rivers were again distinct AUs (Figure 4c, Supplementary Figure 9c). The rivers of southwest England and Brittany, along with the Dee and Eden were a third AU and the remaining British Isles rivers with the Adour constituted the final AU.

## 3.6 Conservation priorities

Results for the expected contribution to diversity measure showed that only 14 and ten rivers made positive contributions to total allelic (*A*_T_) and gene diversity (*H*_T_), respectively (Figure 5a & b, Supplementary Figure 10). For allelic diversity, four non-chalk and all ten chalk rivers had a positive contribution to the metric. For gene diversity, only the Hampshire Basin and Upper Normandy rivers made positive contributions (Figure 5b). At a regional level, the Hampshire Basin rivers had the largest influence on both metrics, especially for total gene diversity, where the contribution was in excess of 6% (Figure 5c & d). The Hampshire Basin populations also had a negative contribution to both within-region metrics (*H*_S_ and *A*_S_; Supplementary Figure 10), suggesting all seven populations have low within-river diversity. The Hampshire Basin and Upper Normandy rivers also made positive contributions to *D*_G_ and *D*_A_ (Supplementary Figure 10) signifying that they contribute substantially to both between-river and between-regional diversity.

## Discussion

Here we present results on the analysis of samples from 42 Atlantic salmon rivers from the British Isles and France with a low-density SNP panel. For the first time, samples from chalk rivers in southern England and northern France are incorporated into the same study. Multiple analyses of the genetic data confirmed the strong differentiation of these English Channel chalk river salmon from those from non-chalk rivers, with outlier tests of SNP data suggesting that this could be due to local adaptation to water chemistry. We identified two potential Evolutionarily Significant Units, with the chalk river salmon also being identified as conservation prioritises based on their unique and substantial contributions to allelic and gene diversity.

### 4.1 Strong regional genetic structuring

The results suggest that there is strong hierarchical structuring of the genetic diversity present within the salmon rivers investigated here, with four main regional genetic groups; English chalk, French chalk, English non-chalk, French non-chalk populations, with further subdivision being present within both English and French non-chalk river populations as observed in many previous studies of Atlantic salmon. Indeed, such hierarchical structuring is a common pattern in Atlantic salmon in which, at the broadest scales, genetic groups can often span hundreds of kilometres of coastline (Bradbury et al., 2014; Cauwelier et al., 2018; Finnegan et al., 2013; Gilbey et al., 2018; Griffiths et al., 2010; Wennevik et al., 2019). Differentiation between rivers within each of the four main groups in the current study was generally low; the average pairwise *F*_ST_ between rivers within each of the four groups were all less than 0.035, indicative of low differentiation (Hartl & Clark, 1997).

Anadromous salmonids often form metapopulations, where rivers are connected by varying levels of migration between them (Lamarins et al., 2024; Schtickzelle & Quinn, 2007). For instance, using otolith chemistry, Fontaine et al. (2025) found frequent dispersal of salmon between three southern French salmon rivers separated by 300 km of coastline, while salmon straying from the River Imsa (Norway) predominately entered rivers within 60 km of the river mouth (Jonsson et al., 2003). Patterns suggestive of frequent dispersal of fish between rivers are also apparent in our results. Salmon from several, geographically proximate rivers showed extremely low levels of differentiation (<0.007; i.e. the Test and Itchen and Tamar and Lynher), indicative of high numbers of straying fish between adjacent rivers. Additionally, there were many cases of pairs of populations within genetic groups demonstrating low differentiation despite being separated by several hundreds of kilometres. For example, the coastal geographic distance between the rivers Exe and the Dee is approximately 700 km, while pairwise *F*_ST_ between the salmon populations in these two rivers was 0.02.

### 4.2 Strong divergence between chalk and non-chalk rivers

In agreement with previous studies utilising microsatellite analysis (Finnegan et al., 2013; Griffiths et al., 2010; Ikediashi et al., 2018; Perrier et al., 2011), SNP markers reaffirmed the uniqueness of the chalk stream salmon populations with multiple analyses highlighting the divergence between fish residing in rivers flowing into the western and eastern parts of the English Channel. For instance, the salmon in southwest England and Brittany in northwest France are more closely related to each other than they are to their nearest neighbours further east along each respective coastline. Interestingly, these patterns in the genetic data correspond precisely with the underlying geology/water chemistry experienced by each population. Brittany and southern Devon/Cornwall are dominated by Devonian age bedrock with granitic intrusions, resulting in more acidic river water (pH≤7) with low conductivity. Additionally, the upland areas of Brittany, Devon and Cornwall are dominated by blanket peat bog, reinforcing the acidic nature of river water in these areas. Further east along both coasts in Normandy and southern England the underlying geology is dominated by Cretaceous era limestones and chalks, resulting in river water with higher pH above 7.

Multiple, interacting factors help determine the chemical composition of river water. Of particular importance is underlying geology which has a strong influence on pH, conductivity and concentrations of dissolved ions (Jarvie et al., 2002; Liu et al., 2000; Rothwell et al., 2010). It has been suggested that the geological characteristics, and therefore chemical characteristics, of river catchments may be an important factor in determining the accuracy of homing through olfactory-based imprinting during smolting (Keefer & Caudill, 2014) and may help to maintain regional population structure via reduced straying of fish between regional groups of rivers (Almodóvar et al., 2023; Bourret et al., 2013).

Additionally, underlying geology and its effect on river water chemistry has been proposed to be a selective agent in the process of local adaptation in Canadian Atlantic salmon and English populations of brown trout (*Salmo trutta*) exposed to high levels of heavy metals (Bourret et al., 2013; Paris et al., 2024). It is interesting to note that the hierarchical genetic structure found here in Atlantic salmon in Channel rivers also occurs in brown trout populations inhabiting rivers flowing into the Channel, with these patterns also having been linked to underlying geology (King et al., 2024). The locations of transitions between groups being coincident in both species reinforces the suggestion that underlying geology and/or water chemistry is playing an important role in driving local adaptation in salmonids residing in rivers entering the English Channel.

Finally, the finding of multiple SNPs under divergent selection adds strength to the assertion that the observed patterns may be driven by local adaptation. The majority of outlier SNPs showed strong allele frequency differences between chalk and non-chalk rivers with some loci e.g. Ssa_58789 and Ssa_24091, being almost completely monomorphic in the non-chalk rivers but polymorphic in chalk rivers. Unfortunately, the strongest candidate SNP (Ssa_69865) is found in a non-coding region on linkage group (LG) ssa13. The closest gene, integrator complex subunit 2 (ints2), is approximately 48 000 base pairs upstream of the Ssa_69865 polymorphism.

We discovered that a SNP linked to the gene SIX homeobox 6 (six6) was under divergent section in the full dataset and non-chalk-only outlier tests, with the region containing this polymorphism having been shown to be under selection across both the North American and European ranges of Atlantic salmon (Barson et al., 2015; Cauwelier et al., 2018; Kess et al., 2024; V. L. Pritchard et al., 2018; Zueva et al., 2021). This region, on LG ssa09, has been implicated in variation in adult run timing in Scottish salmon populations (Cauwelier et al., 2018) and age at maturity in Scandinavian salmon (Barson et al., 2015) and suggests that there may be differences in these two life history metrics across the range of the populations sampled here. For instance, the rivers of northeast England tend to have a higher proportion of multi-sea winter (MSW) salmon in their rod catches than the rivers of southern and southwest England (Environment Agency, 2024). Likewise, the rivers of southern France, including the Adour, are dominated by MSW salmon compared to the rivers of Normandy and Brittany (Bal et al., 2017; Le Cam et al., 2015). The highest MAF for Ssa_six6 were found in the rivers of the Bristol Channel, a major inlet separating south Wales from southwest England. The rivers of the inner Bristol Channel (Usk, Wye and Severn) are atypical from the majority of the rivers studied here, being large catchments where adult salmon must undertake extensive freshwater migrations (>100-300 km) to reach spawning grounds. Indeed, the rod catch of these rivers are dominated by large multi-sea winter salmon (Environment Agency, 2024). Further highlighting the significance of this genomic region, six6 has also been found to be an important determinant of age at maturity in Pacific salmonids (Waters et al., 2021; Willis et al., 2020). With the exception of six6, based on the small number of loci screened here we cannot yet identify any obvious relationship between the outlier loci and any biological function that could account for the distinction between chalk and non-chalk salmon. However, the outlier loci are spread across multiple chromosomes and possibly indicate that the differences between chalk and non-chalk salmon are polygenic in nature. Additional investigation, using reduced representation or whole genome sequencing, will be required to explore the molecular mechanisms underlying local adaptation in chalk salmon.

### 4.3 Low divergence within genetic clusters

Anadromous species, such as salmon and trout, are known for their ability to home to their natal river after spending time feeding at sea. This spawning philopatry has profound evolutionary consequences, often leading to reduced gene flow between populations and giving rise to the within-river genetic structuring often observed in salmonid species (Dillane et al., 2008; Pritchard et al., 2018). However, straying of fish to non-natal rivers does also occur (Griffiths et al., 2011; Perrier et al., 2011) and is recognised as an important evolutionary feature of salmonids, buffering populations against spatial and temporal variation in local environments (Keefer & Caudill, 2014).

Multiple analyses suggest relatively low levels of divergence between salmon populations within each of the four main groups of rivers. Pairwise *F*_ST_ values indicate very low levels of differentiation, especially between geographically proximate rivers such as the Frome and Piddle (F_ST_=0.006) and the Seé and Sienne (F_ST_=-0.003). Moreover, while DAPC and STRUCTURE analyses suggest that there may be some genetic structuring within the Hampshire Basin chalk rivers, there was only a limited geographical component to this diversity (a relatively weak split between salmon populations in the eastern and western Hampshire Basin rivers).

The freshwater homing of salmon is driven primarily by olfactory imprinting of natal waters, with parr/smolts ‘learning’ the chemical cues of their natal stream prior to seaward migration (Hasler et al., 1978). Increased straying between rivers with similar water chemistry profiles has been found for Canadian Atlantic salmon populations (Bradbury et al., 2014). Given that all the Hampshire Basin rivers originate from the same chalk aquifer system (Allen, 2017), the low divergence between salmon populations in these rivers may reflect a high degree of chemical similarity of the water in each of the Hampshire Basin rivers (at least with regard to the homing cues recognised by Atlantic salmon).

### 4.4 Genetic signal of historic stocking in Upper Normandy populations

The DAPC and STRUCTURE analyses showed clearly that two individuals sampled from two French chalk rivers assigned genetically to non-chalk rivers. These individuals were sampled as adults and two plausible explanations can be advanced to explain this observation. The first is that the two fish in question are strays from non-chalk rivers. Straying is a common phenomenon in anadromous salmonids, facilitating colonisation of new habitats and gene flow between rivers (Lamarins et al., 2024), with well-documented examples of both short- and long-distance movements of salmon between rivers within our study area having been reported previously (Griffiths et al., 2011; Leunda et al., 2013; Perrier et al., 2010). Alternatively, these fish might constitute the remnants of historical stocking programmes. Upper Normandy rivers experienced ‘high’ stocking intensity up to 1990, predominately with salmon from rivers on the east coast of Scotland (Perrier et al., 2013). Previous microsatellite-based analysis suggested that the genetic effects of this stocking were short-lived in these rivers (Perrier et al., 2013). In contrast, however, our SNP data suggest a longer-lasting impact of stocking on genetic diversity within Upper Normandy salmon, with the rivers of this region having the highest values for basic measures of diversity (H_O_ & H_S_) calculated in the current study and the STRUCTURE analysis showing a significant contribution of non-chalk ancestry.

### 4.5 Salmon Conservation Units

Genetic and genomic data have been used to delimit Conservation Units (ESUs, MUs and AUs) in salmonids that might represent important components of diversity in need of specific conservation and management measures (Lehnert et al., 2023; Waples, 1991; Xuereb et al., 2022). A broad definition of an ESU is a group of populations that merit separate management and conservation as a consequence of genetic and ecological distinctiveness (Allendorf et al., 2022). Some definitions also include a degree of geographic isolation (Funk et al., 2012). Our results indicate that the Atlantic salmon rivers studied here could be conservatively classified into two distinct Evolutionarily Significant Units (chalk and non-chalk) on the basis of genetic, ecological and geographic distinctiveness. Each of these two ESUs can be further subdivided into multiple distinct MUs. However, to accurately delimit management units, a more exhaustive sampling of populations in the study area, particularly for northern France, is required. Similarly, given the small number of outlier loci identified here, a more extensive genomic analysis will be needed to accurately define the adaptive units across the regions investigated here.

### 4.6 Chalk salmon as a distinct sub-species

The criteria that are used to distinguish ESUs are similar to those used to delineate a subspecies. Taylor et al. (2017) defined a subspecies as ‘a population, or collection of populations, that appears to be a separately evolving lineage with discontinuities resulting from geography, ecological specialization, or other forces that restrict gene flow to the point that the population or collection of populations is diagnosably distinct’.

The chalk stream salmon fulfil all three requirements. They are a genetically distinct and definable unit found in a geographically restricted area. Critically, the rivers they inhabit have well-defined consistent geochemical, morphological and hydrodynamic characteristics (namely all the criteria which define a chalk stream). They (the chalk stream salmon - or, more precisely, fish with their genotypes and allelic profiles) are not found anywhere outside of a well-defined geographic region. Assignment to regions is robust and evidence of straying is virtually absent; in the English chalk stream populations there is no evidence of any successful straying leading to gene flow between chalk and non-chalk populations.

Taken into consideration, we suggest that the Atlantic salmon populations of the chalk streams meet the criteria for recognition as a subspecies within the species *Salmo salar* L. and suggest the taxonomic name *Salmo salar calcariensis* for this ESU. However, more extensive genomic studies are needed to fully address the genomic basis of intraspecific taxonomy of salmon in this region

### 4.7 The conservation value of chalk stream salmon

It is clear that concerted conservation action is required to halt the continuing decline in both the numbers and genetic diversity observed across the species’ range of Atlantic salmon (Lehnert et al., 2019). Moreover, as different environmental stressors act across different parts of the species’ range, the continuing preservation of healthy regional populations is a key concern for conservation management of the species (von Takach et al., 2024).

In particular, the UK chalk stream salmon populations are highlighted for their unique genetic make-up and the importance of their contribution to the overall diversity of the species. All had negative contributions to both H_S_ and A_S_ (Supplementary Figure 8), suggesting the populations in the seven studied rivers have low within-river diversity. This is re-enforced by these populations generally having the lowest observed and expected heterozygosities and locus polymorphism of the populations studied. The significant positive contributions to between-population metrics and negative within-population contributions suggest that these rivers have been affected by long-term isolation, genetic drift and/or bottlenecks (von Takach et al., 2024). Given that the populations in the Hampshire Basin rivers are in continued declining (Environment Agency, 2024) and that some rivers have experienced significantly poor recruitment in recent years (UK Environment Agency & GWCT, personal communications), there is need for enhanced conservation and protection above that already afforded under United Kingdom law.

### 4.8 Conclusions

In summary, we propose that the chalk stream salmon of southern England and northern France constitute a unique component of the Atlantic salmon meta-population; they contain a distinctive and substantial component of the genetic diversity of the species and this degree of genetic novelty is such that they constitute a distinct ESU, linked primarily to the underlying geology and water chemistry of their local habitat. As such, their extirpation would result in the loss of a significant component of the overall genetic diversity (>6%) of the species (based on the populations included in the current study) and we reiterate the urgent need to preserve them and the rivers in which they reside.

Genetically distinct populations should be prioritised for conservation as these often represent unique evolutionary lineages, within the overall metapopulation structure of species such as Atlantic salmon. Indeed, one of the overarching strengths of conservation genetics is the ability to define evolutionarily divergent units (populations) within a species that warrant recognition and separate/bespoke management. Chalk salmon appear to represent such a unit and may be potentially viewed either as a distinct ESU or as a new subspecies, *Salmo salar calcariensis*.

## Acknowledgements

We thank Simon Toms and the staff of the Environment Agency, Celine Artero, Daniel Osmond and Charlie Ellis for co-ordinating and supporting sample collection. The French adult scale samples used in this study were provided by the Biological Resource Centre Colisa (https://doi.org/10.15468/q99pu9) part of BRC4Env (DOI: https://doi.org/10.15454/TRBJTB), of the Research Infrastructure AgroBRC-RARe.

## Funding Information

This research was funded by the European Union Interreg France (Channel) England programme project ‘Salmonid Management Around the Channel’ (SAMARCH) with additional funding from Southern Water, UK.

## Notes

### Competing Interest Statement

The authors have declared no competing interest.

## References

Allen, D. J. (2017). The chalk aquifer of the Wessex Basin. British Geological Survey Research Report No. RR/11/02.

Allendorf, F. W., Funk, W. C., Aitken, S. N., Byrne, M., & Luikart, G. (2022). Conservation and the Genomics of Populations (Third edition ed.). Oxford, UK: Oxford University Press.

Almodóvar, A., Nicola, G. G., Ayllón, D., Leal, S., Marchán, D. F., & Elvira, B. (2023). A benchmark for Atlantic salmon conservation: genetic diversity and structure in a southern European glacial refuge before the climate changed. Fishes, 8(6), 321. doi:10.3390/fishes8060321

Anderson, E. C., Clemento, A. J., Campell, M. A., Pearse, D. E., Beulke, A. K., Columbus, C., Garza, J. C. (2025). A multipurpose microhaplotype panel for genetic analysis of California Chinook salmon. Evolutionary Applications, 18(5), e70110. doi:10.1111/eva.70110

Bal, G., Montorio, L., Rivot, E., Prévost, E., Baglinière, J. L., & Nevoux, M. (2017). Evidence for long-term change in length, mass and migration phenology of anadromous spawners in French Atlantic salmon Salmo salar. Journal of Fish Biology, 90(6), 2375–2393. doi:10.1111/jfb.13314

Barson, N. J., Aykanat, T., Hindar, K., Baranski, M., Bolstad, G. H., Fiske, P., Primmer, C. R. (2015). Sex-dependent dominance at a single locus maintains variation in age at maturity in salmon. Nature, 528(7582), 405–+. doi:10.1038/nature16062

Beaumont, M. A., & Balding, D. J. (2004). Identifying adaptive genetic divergence among populations from genome scans. Molecular Ecology, 13(4), 969–980. doi:10.1111/j.1365-294X.2004.02125.x

Berrie, A. D. (1992). The chalk-stream environment. Hydrobiologia, 248(1), 3–9. doi:10.1007/bf00008881

Bourret, V., Dionne, M., Kent, M. P., Lien, S., & Bernatchez, L. (2013). Landscape genomics in Atlantic salmon (Salmo salar): searching for gene-environment interactions driving local adaptation. Evolution, 67(12), 3469–3487. doi:10.1111/evo.12139

Bradbury, I. R., Hamilton, L. C., Robertson, M. J., Bourgeois, C. E., Mansour, A., & Dempson, J. B. (2014). Landscape structure and climatic variation determine Atlantic salmon genetic connectivity in the Northwest Atlantic. Canadian Journal of Fisheries and Aquatic Sciences, 71(2), 246–258. doi:10.1139/cjfas-2013-0240

Bradbury, I. R., Lehnert, S. J., Messmer, A., Duffy, S. J., Verspoor, E., Kess, T., Reddin, D. (2021). Range-wide genetic assignment confirms long-distance oceanic migration in Atlantic salmon over half a century. Ices Journal of Marine Science, 78(4), 1434–1443. doi:10.1093/icesjms/fsaa152

CABA. (2021). Chalk stream restoration strategy 2021. Retrieved from https://catchmentbasedapproach.org/wp-content/uploads/2022/11/CaBA-CSRG-IMP-PLAN-FINAL-25.11.22.-V2.pdf

Caballero, A., & Rodríguez-Ramilo, S. T. (2010). A new method for the partition of allelic diversity within and between subpopulations. Conservation Genetics, 11(6), 2219–2229. doi:10.1007/s10592-010-0107-7

Cauwelier, E., Gilbey, J., Sampayo, J., Stradmeyer, L., & Middlemas, S. J. (2018). Identification of a single genomic region associated with seasonal river return timing in adult Scottish Atlantic salmon (Salmo salar), using a genome-wide association study. Canadian Journal of Fisheries and Aquatic Sciences, 75(9), 1427–1435. doi:10.1139/cjfas-2017-0293

Cauwelier, E., Verspoor, E., Coulson, M. W., Armstrong, A., Knox, D., Stradmeyer, L., Gilbey, J. (2018). Ice sheets and genetics: Insights into the phylogeography of Scottish Atlantic salmon, Salmo salar L. Journal of Biogeography, 45(1), 51–63. doi:10.1111/jbi.13097

Cavalli-Sforza, L. L., & Edwards, A. W. F. (1967). Phylogenetic analysis models and estimation procedures. American Journal of Human Genetics, 19(3P1), 233–257.

Chhina, A. K., Abhari, N., Mooers, A., & Lewthwaite, J. M. M. (2024). Linking the spatial and genomic structure of adaptive potential for conservation management: a review. Genome, 21. doi:10.1139/gen-2024-0036

Comte, L., Buisson, L., Daufresne, M., & Grenouillet, G. (2013). Climate-induced changes in the distribution of freshwater fish: observed and predicted trends. Freshwater Biology, 58(4), 625–639. doi:10.1111/fwb.12081

Crisp, D. T. (2000). Trout and SAlmon: Ecolopgy, Conservation and Rehabilitation. Oxford: Fishing News Books/Balckwell Science.

Dadswell, M., Spares, A., Reader, J., McLean, M., McDermott, T., Samways, K., & Lilly, J. (2022). The decline and impending collapse of the Atlantic salmon (Salmo salar) population in the North Atlantic ocean: a review of possible causes. Reviews in Fisheries Science & Aquaculture, 30(2), 215–258. doi:10.1080/23308249.2021.1937044

Darwell, W. R. T. (2023a). Salmo salar (English Chalkstream subpopulation). The IUCN Red List of Threatened Species 2023: e.T196579553A196580841. Retrieved from 10.2305/IUCN.UK.2023-1.RLTS.T213546282A213546288.en

Darwell, W. R. T. (2023b). Salmo salar (France/Spain subpopulation). The IUCN Red List of Threatened Species 2023: e.T196579750A196580901. Retrieved from 10.2305/IUCN.UK.2023-1.RLTS.T196579750A196580901.en.

Darwell, W. R. T. (2023c). Salmo salar (Ireland subpopulation). The IUCN Red List of Threatened Species 2023: e.T214416825A214416828. Retrieved from 10.2305/IUCN.UK.2023-1.RLTS.T214416825A214416828.en.

Darwell, W. R. T. (2023d). Salmo salar. The IUCN Red List of Threatened Species 2023: e.T19855A67373433. Retrieved from 10.2305/IUCN.UK.2023-1.RLTS.T19855A67373433.en

Deinet, S., Flint, R., Puleston, H., Baratech, A., Royte, J., Thieme, M. L., Wanningen, H. (2024). The Living Planet Index (LPI) for migratory freshwater fish 2024 update Technical Report. Retrieved from The Netherlands:

Dillane, E., McGinnity, P., Coughlan, J. P., Cross, M. C., de Eyto, E., Kenchington, E., Cross, T. F. (2008). Demographics and landscape features determine intrariver population structure in Atlantic salmon (Salmo salar L.): the case of the River Moy in Ireland. Molecular Ecology, 17(22), 4786–4800. doi:10.1111/j.1365-294X.2008.03939.x

Dudgeon, D. (2019). Multiple threats imperil freshwater biodiversity in the Anthropocene. Current Biology, 29(19), R960–R967. doi:10.1016/j.cub.2019.08.002

Environment Agency, E. (2024). Salmonid and fisheries statistics for England and Wales 2023. Retrieved from https://www.gov.uk/government/publications/assessment-of-salmon-stocks-and-fisheries-england-and-wales-2023

Evanno, G., Legrand, M., Nevoux, M., Prevost, E., & Phillips, K. (2023). Salmo salar (Allier subpopulation). The IUCN Red List of Threatened Species 2023: e.T229537556A229549238. Retrieved from 10.2305/IUCN.UK.2023-1.RLTS.T229537556A229549238.en

Evanno, G., Regnaut, S., & Goudet, J. (2005). Detecting the number of clusters of individuals using the software STRUCTURE: a simulation study. Molecular Ecology, 14(8), 2611–2620. doi:10.1111/j.1365-294X.2005.02553.x

Excoffier, L., Hofer, T., & Foll, M. (2009). Detecting loci under selection in a hierarchically structured population. Heredity, 103, 285–298. doi:10.1038/hdy.2009.74

Excoffier, L., & Lischer, H. E. L. (2010). Arlequin suite ver 3.5: a new series of programs to perform population genetics analyses under Linux and Windows. Molecular Ecology Resources, 10(3), 564–567. doi:10.1111/j.1755-0998.2010.02847.x

Exposito-Alonso, M., Booker, T. R., Czech, L., Gillespie, L., Hateley, S., Kyriazis, C. C., Zess, E. (2022). Genetic diversity loss in the Anthropocene. Science, 377(6613), 1431–1435. doi:10.1126/science.abn5642

Fick, S. E., & Hijmans, R. J. (2017). WorldClim 2: new 1-km spatial resolution climate surfaces for global land areas. International Journal of Climatology, 37(12), 4302–4315. doi:10.1002/joc.5086

Finnegan, A. K., Griffiths, A. M., King, R. A., Machado-Schiaffino, G., Porcher, J. P., Garcia-Vazquez, E., Stevens, J. R. (2013). Use of multiple markers demonstrates a cryptic western refugium and postglacial colonisation routes of Atlantic salmon (Salmo salar L.) in northwest Europe. Heredity, 111(1), 34–43. doi:10.1038/hdy.2013.17

Foll, M., & Gaggiotti, O. (2008). A genome-scan method to identify selected loci appropriate for both dominant and codominant markers: a Bayesian perspective. Genetics, 180(2), 977–993. doi:10.1534/genetics.108.092221

Fontaine, A., Vignon, M., Tabouret, H., Holub, A., Barranco, G., Bosc, S., Bareille, G. (2025). Inter-annual dispersal stability within the Atlantic salmon metapopulation from the Bay of Biscay. Fisheries Research, 284, 107323. doi:10.1016/j.fishres.2025.107323

Forester, B. R., Lasky, J. R., Wagner, H. H., & Urban, D. L. (2018). Comparing methods for detecting multilocus adaptation with multivariate genotype-environment associations. Molecular Ecology, 27(9), 2215–2233. doi:10.1111/mec.14584

Forrester, B. R., & Lama, T. M. (2023). The role of genomics in the future of ESA decision-making. In The Codex of the Endangered Species Act, Volume II: The Next Fifty Years (pp 159–186). Lanham: Rowman & Littlefield Publishers.

Francis, R. M. (2017). POPHELPER: an R package and web app to analyse and visualize population structure. Molecular Ecology Resources, 17(1), 27–32. doi:10.1111/1755-0998.12509

Funk, W. C., McKay, J. K., Hohenlohe, P. A., & Allendorf, F. W. (2012). Harnessing genomics for delineating conservation units. Trends in Ecology & Evolution, 27(9), 489–496. doi:10.1016/j.tree.2012.05.012

Gilbey, J., Coughlan, J., Wennevik, V., Prodöhl, P., Stevens, J. R., de Leaniz, C. G., Verspoor, E. (2018). A microsatellite baseline for genetic stock identification of European Atlantic salmon (Salmo salar L.). Ices Journal of Marine Science, 75(2), 662–674. doi:10.1093/icesjms/fsx184

Gillson, J. P., Bašić, T., Davison, P., Riley, W. D., Talks, L., Walker, A. M., & Russell, I. C. (2022). A review of marine stressors impacting Atlantic salmon Salmo salar, with an assessment of the major threats to English stocks. Reviews in Fish Biology and Fisheries, 32(3), 879–919. doi:10.1007/s11160-022-09714-x

Goldberg, C. S., & Waits, L. P. (2010). Quantification and reduction of bias from sampling larvae to infer population and landscape genetic structure. Molecular Ecology Resources, 10(2), 304–313. doi:10.1111/j.1755-0998.2009.02755.x

Griffiths, A. M., Ellis, J. S., Clifton-Dey, D., Machado-Schiaffino, G., Bright, D., Garcia-Vazquez, E., & Stevens, J. R. (2011). Restoration versus recolonisation: The origin of Atlantic salmon (Salmo salar L.) currently in the River Thames. Biological Conservation, 144(11), 2733–2738. doi:10.1016/j.biocon.2011.07.017

Griffiths, A. M., Machado-Schiaffino, G., Dillane, E., Coughlan, J., Horreo, J. L., Bowkett, A. E., Stevens, J. R. (2010). Genetic stock identification of Atlantic salmon (Salmo salar) populations in the southern part of the European range. Bmc Genetics, 11, 27. doi:10.1186/1471-2156-11-31

Haase, P., Cortés-Guzmán, D., He, F., Jupke, J. F., Mangadze, T., Pelicice, F. M., Sinclair, J. S. (2025). Successes and failures of conservation actions to halt global river biodiversity loss. Nature Reviews Biodiversity. doi:10.1038/s44358-024-00012-x

Hansen, M. M., Nielsen, E. E., & Mensberg, K. L. D. (1997). The problem of sampling families rather than populations: Relatedness among individuals in samples of juvenile brown trout Salmo trutta L. Molecular Ecology, 6(5), 469–474. doi:10.1046/j.1365-294X.1997.t01-1-00202.x

Hartl, D. L., & Clark, A. G. (1997). Principles of Population Genetics (3rd Edition ed.). Sunderland, MA: Sinauer Associates, Inc.

Hasler, A. D., Scholz, A. T., & Horrall, R. M. (1978). Olfactory imprinting and homing in salmon. American Scientist, 66(3), 347–355.

Hijmans, R. J. (2019). Geographic data analysis and modelling. R package version 3. 0-7. Retrieved from https://cran.r-project.org/web/packages/raster/index.html

Ikediashi, C., Paris, J. R., King, R. A., Beaumont, W. R. C., Ibbotson, A., & Stevens, J. R. (2018). Atlantic salmon Salmo salar in the chalk streams of England are genetically unique. Journal of Fish Biology, 92(3), 621–641. doi:10.1111/jfb.13538

Jarvie, H. P., Oguchi, T., & Neal, C. (2002). Exploring the linkages between river water chemistry and watershed characteristics using GIS-based catchment and locality analyses. Regional Environmental Change, 3(1-3), 36–50. doi:10.1007/s10113-001-0036-6

Jeffery, N. W., Wringe, B. F., McBride, M. C., Hamilton, L. C., Stanley, R. R. E., Bernatchez, L., Bradbury, I. R. (2018). Range-wide regional assignment of Atlantic salmon (Salmo salar) using Check tor genome wide single-nucleotide polymorphisms. Fisheries Research, 206, 163–175. doi:10.1016/j.fishres.2018.05.017

Jombart, T. (2008). adegenet: a R package for the multivariate analysis of genetic markers. Bioinformatics, 24(11), 1403–1405. doi:10.1093/bioinformatics/btn129

Jombart, T., Devillard, S., & Balloux, F. (2010). Discriminant analysis of principal components: a new method for the analysis of genetically structured populations. Bmc Genetics, 11, 15. doi:10.1186/1471-2156-11-94

Jones, O. R., & Wang, J. L. (2010). COLONY: a program for parentage and sibship inference from multilocus genotype data. Molecular Ecology Resources, 10(3), 551–555. doi:10.1111/j.1755-0998.2009.02787.x

Jonsson, B., Jonsson, N., & Hansen, L. P. (2003). Atlantic salmon straying from the River Imsa. Journal of Fish Biology, 62(3), 641–657. doi:10.1046/j.1095-8649.2003.00053.x

Keefer, M. L., & Caudill, C. C. (2014). Homing and straying by anadromous salmonids: a review of mechanisms and rates. Reviews in Fish Biology and Fisheries, 24(1), 333–368. doi:10.1007/s11160-013-9334-6

Kess, T., Lehnert, S. J., Bentzen, P., Duffy, S., Messmer, A., Dempson, J. B., Bradbury, I. R. (2024). Variable parallelism in the genomic basis of age at maturity across spatial scales in Atlantic Salmon. Ecology and Evolution, 14(4), 18. doi:10.1002/ece3.11068

King, R. A., Ellis, C. D., Bekkevold, D., Ensing, D., Lecointre, T., Osmond, D. R., Stevens, J. R. (2024). Leveraging the genetic diversity of trout in the rivers of the British Isles and northern France to understand the movements of sea trout (Salmo trutta L.) around the English Channel. Evolutionary Applications, 17(7), 15. doi:10.1111/eva.13759

King, R. A., & Stevens, J. R. (2021). Development of SNP markers derived from RAD sequencing for Atlantic salmon (Salmo salar L.) inhabiting the rivers of southern England. Conservation Genetics Resources, 13(4), 369–373. doi:10.1007/s12686-021-01215-6

Klemetsen, A., Amundsen, P. A., Dempson, J. B., Jonsson, B., Jonsson, N., O’Connell, M. F., & Mortensen, E. (2003). Atlantic salmon Salmo salar L., brown trout Salmo trutta L. and Arctic charr Salvelinus alpinus (L.):: a review of aspects of their life histories. Ecology of Freshwater Fish, 12(1), 1–59. doi:10.1034/j.1600-0633.2003.00010.x

Lamarins, A., Carlson, S. M., & Buoro, M. (2024). Dispersal and gene flow in anadromous salmonids: A systematic review. Ecology of Freshwater Fish, 33(4), e12811. doi:10.1111/eff.12811

Langella, O. (1999). Populations v1.2.28. Retrieved from http://bioinformatics.org/populations/

Le Cam, S., Perrier, C., Besnard, A. L., Bernatchez, L., & Evanno, G. (2015). Genetic and phenotypic changes in an Atlantic salmon population supplemented with non-local individuals: a longitudinal study over 21 years. Proceedings of the Royal Society B-Biological Sciences, 282(1802), 9. doi:10.1098/rspb.2014.2765

Lehnert, S. J., Bradbury, I. R., Wringe, B. F., Van Wyngaarden, M., & Bentzen, P. (2023). Multifaceted framework for defining conservation units: An example from Atlantic salmon (Salmo salar) in Canada. Evolutionary Applications, 16(9), 1568–1585. doi:10.1111/eva.13587

Lehnert, S. J., Kess, T., Bentzen, P., Kent, M. P., Lien, S., Gilbey, J., Bradbury, I. R. (2019). Genomic signatures and correlates of widespread population declines in salmon. Nature Communications, 10, 10. doi:10.1038/s41467-019-10972-w

Leunda, P. M., Ardaiz, J., Russell, I. C., Toms, S., & Hillman, R. (2013). Homing and straying of Atlantic salmon in the Bidasoa River: report of an unusual stray from Great Britain to the Iberian Peninsula. Fisheries Management and Ecology, 20(5), 460–463. doi:10.1111/fme.12029

Liu, Z. J., Weller, D. E., Correll, D. L., & Jordan, T. E. (2000). Effects of land cover and geology on stream chemistry in watersheds of Chesapeake Bay. Journal of the American Water Resources Association, 36(6), 1349–1365. doi:10.1111/j.1752-1688.2000.tb05731.x

López-Cortegano, E., Pérez-Figueroa, A., & Caballero, A. (2019). metapop2: Re-implementation of software for the analysis and management of subdivided populations using gene and allelic diversity. Molecular Ecology Resources, 19(4), 1095–1100. doi:10.1111/1755-0998.13015

Meirmans, P. G. (2020). GENODIVE version 3.0: Easy-to-use software for the analysis of genetic data of diploids and polyploids. Molecular Ecology Resources, 20(4), 1126–1131. doi:10.1111/1755-0998.13145

Menyhart, O., Weltz, B., & Gyorffy, B. (2021). MultipleTesting.com: A tool for life science researchers for multiple hypothesis testing correction. Plos One, 16(6), 12. doi:10.1371/journal.pone.0245824

Nei, N. (1987). Molecular Evolutionary Genetics. New York: Columbia University Press.

Nunn, A. D., Ainsworth, R. F., Walton, S., Bean, C. W., Hatton-Ellis, T. W., Brown, A., Noble, R. A. A. (2023). Extinction risks and threats facing the freshwater fishes of Britain. Aquatic Conservation-Marine and Freshwater Ecosystems, 33(12), 1460–1476. doi:10.1002/aqc.4014

O’Sullivan, R. J., Ozerov, M., Bolstad, G. H., Gilbey, J., Jacobsen, J. A., Erkinaro, J., Aykanat, T. (2022). Genetic stock identification reveals greater use of an oceanic feeding ground around the Faroe Islands by multi-sea winter Atlantic salmon, with variation in use across reporting groups. Ices Journal of Marine Science, 79(9), 2442–2452. doi:10.1093/icesjms/fsac182

Oksanen, J., Guillaume Blanchet, F., Friendly, M., Kindt, R., Legendre, P., McGlinn, D., Wagner, H. (2019). Vegan: Community Ecology Package. R Package Version 2. 5-2. Retrieved from https://cran.r-project.org/web/packages/vegan/vegan.pdf.

Pante, E., & Simon-Bouhet, B. (2013). marmap: A package for importing, plotting and analyzing bathymetric and topographic data in R. Plos One, 8(9), 4. doi:10.1371/journal.pone.0073051

Paris, J. R., King, R. A., Obiol, J. F., Shaw, S., Lange, A., Bourret, V., Stevens, J. R. (2024). The Genomic Signature and Transcriptional Response of Metal Tolerance in Brown Trout Inhabiting Metal-Polluted Rivers. Molecular Ecology, 19. doi:10.1111/mec.17591

Perrier, C., Baglinière, J. L., & Evanno, G. (2013). Understanding admixture patterns in supplemented populations: a case study combining molecular analyses and temporally explicit simulations in Atlantic salmon. Evolutionary Applications, 6(2), 218–230. doi:10.1111/j.1752-4571.2012.00280.x

Perrier, C., Evanno, G., Belliard, J., Guyomard, R., & Baglinière, J. L. (2010). Natural recolonization of the Seine River by Atlantic salmon (Salmo salar) of multiple origins. Canadian Journal of Fisheries and Aquatic Sciences, 67(1), 1–4. doi:10.1139/f09-190

Perrier, C., Guyomard, R., Bagliniere, J. L., & Evanno, G. (2011). Determinants of hierarchical genetic structure in Atlantic salmon populations: environmental factors vs anthropogenic influences. Molecular Ecology, 20(20), 4231–4245. doi:10.1111/j.1365-294X.2011.05266.x

Petit, R. J., El Mousadik, A., & Pons, O. (1998). Identifying populations for conservation on the basis of genetic markers. Conservation Biology, 12(4), 844–855. doi:10.1046/j.1523-1739.1998.96489.x

Pritchard, J. K., Stephens, M., & Donnelly, P. (2000). Inference of population structure using multilocus genotype data. Genetics, 155(2), 945–959.

Pritchard, V. L., Mäkinen, H., Vähä, J. P., Erkinaro, J., Orell, P., & Primmer, C. R. (2018). Genomic signatures of fine-scale local selection in Atlantic salmon suggest involvement of sexual maturation, energy homeostasis and immune defence-related genes. Molecular Ecology, 27(11), 2560–2575. doi:10.1111/mec.14705

Raymond, M., & Rousset, F. (1995). GENEPOP (version-1.2) - population genetics software for exact tests and ecumenicism. Journal of Heredity, 86(3), 248–249. doi:10.1093/oxfordjournals.jhered.a111573

Rodríguez-Ramilo, S. T., & Wang, J. L. (2012). The effect of close relatives on unsupervised Bayesian clustering algorithms in population genetic structure analysis. Molecular Ecology Resources, 12(5), 873–884. doi:10.1111/j.1755-0998.2012.03156.x

Rothwell, J. J., Dise, N. B., Taylor, K. G., Allott, T. E. H., Scholefield, P., Davies, H., & Neal, C. (2010). A spatial and seasonal assessment of river water chemistry across North West England. Science of the Total Environment, 408(4), 841–855. doi:10.1016/j.scitotenv.2009.10.041

Samy, J. K. A., Mulugeta, T. D., Nome, T., Sandve, S. R., Grammes, F., Kent, M. P., Våge, D. I. (2017). SalmoBase: an integrated molecular data resource for Salmonid species. Bmc Genomics, 18, 5. doi:10.1186/s12864-017-3877-1

Schtickzelle, N., & Quinn, T. P. (2007). A metapopulation perspective for salmon and other anadromous fish. Fish and Fisheries, 8(4), 297–314. doi:10.1111/j.1467-2979.2007.00256.x

Sear, D. A., Armitage, P. D., & Dawson, F. H. (1999). Groundwater dominated rivers. Hydrological Processes, 13(3), 255–276. doi:10.1002/(sici)1099-1085(19990228)13:3<255::Aid-hyp737>3.0.Co;2-y

Shaw, R. E., Farquharson, K. A., Bruford, M. W., Coates, D. J., Elliott, C. P., Mergeay, J., Grueber, C. E. (2025). Global meta-analysis shows action is needed to halt genetic diversity loss. Nature, 24. doi:10.1038/s41586-024-08458-x

Tamura, K., Stecher, G., Peterson, D., Filipski, A., & Kumar, S. (2013). MEGA6: Molecular Evolutionary Genetics Analysis Version 6 0. Molecular Biology and Evolution, 30(12), 2725–2729. doi:10.1093/molbev/mst197

Taylor, B. L., Perrin, W. F., Reeves, R. R., Rosel, P. E., Wang, J. Y., Cipriano, F., Brownell, R. L. (2017). Why we should develop guidelines and quantitative standards for using genetic data to delimit subspecies for data-poor organisms like cetaceans. Marine Mammal Science, 33, 12–26. doi:10.1111/mms.12413

Truett, G. E., Heeger, P., Mynatt, R. L., Truett, A. A., Walker, J. A., & Warman, M. L. (2000). Preparation of PCR-quality mouse genomic DNA with hot sodium hydroxide and tris (HotSHOT). Biotechniques, 29(1), 52–+. doi:10.2144/00291bm09

Urban, M. C. (2024). Climate change extinctions. Science, 386, 1123 1128.

von Takach, B., Cameron, S. F., Cremona, T., Eldridge, M. D. B., Fisher, D. O., Hohnen, R., Banks, S. C. (2024). Conservation prioritisation of genomic diversity to inform management of a declining mammal species. Biological Conservation, 291, 12. doi:10.1016/j.biocon.2024.110467

Waples, R. S. (1991). Pacific salmon, Oncorhynchus spp., and the definition of “species” under the Endagered Species Act. Marine Fisheries Review, 53(3), 11–22.

Waples, R. S., & Anderson, E. C. (2017). Purging putative siblings from population genetic data sets: a cautionary view. Molecular Ecology, 26(5), 1211–1224. doi:10.1111/mec.14022

Waters, C. D., Clemento, A., Aykanat, T., Garza, J. C., Naish, K. A., Narum, S., & Primmer, C. R. (2021). Heterogeneous genetic basis of age at maturity in salmonid fishes. Molecular Ecology, 30(6), 1435–1456. doi:10.1111/mec.15822

Weir, B. S., & Cockerham, C. C. (1984). Estimating F-statisticsfor the analysis of popualtions structure. Evolution, 38(6), 1358–1370. doi:10.2307/2408641

Wennevik, V., Quintela, M., Skaala, O., Verspoor, E., Prusov, S., & Glover, K. A. (2019). Population genetic analysis reveals a geographically limited transition zone between two genetically distinct Atlantic salmon lineages in Norway. Ecology and Evolution, 9(12), 6901–6921. doi:10.1002/ece3.5258

Willis, S. C., Hess, J. E., Fryer, J. K., Whiteaker, J. M., Brun, C., Gerstenberger, R., & Narum, S. R. (2020). Steelhead (Oncorhynchus mykiss) lineages and sexes show variable patterns of association of adult migration timing and age-at-maturity traits with two genomic regions. Evolutionary Applications, 13(10), 2836–2856. doi:10.1111/eva.13088

WWF. (2024). Living Planet Report 2024 - A System in Peril. Retrieved from Gland, Switzerland:

WWF-UK. (2015). The State of England’s Chalk Streams. Retrieved from

Xuereb, A., Rougemont, Q., Dallaire, X., Moore, J. S., Normandeau, E., Bougas, B., Bernatchez, L. (2022). Re-evaluating Coho salmon (Oncorhynchus kisutch) conservation units in Canada using genomic data. Evolutionary Applications, 15(11), 1925–1944. doi:10.1111/eva.13489

Zueva, K. J., Lumme, J., Veselov, A. E., Primmer, C. R., & Pritchard, V. L. (2021). Population genomics reveals repeated signals of adaptive divergence in the Atlantic salmon of north-eastern Europe. Journal of Evolutionary Biology, 34(6), 866–878. doi:10.1111/jeb.13732

