## Supplementary for "The chalk streams of southern England and northern France harbour substantial unique components of the overall genetic diversity of Atlantic salmon (*Salmo salar* L.)"

**Supplementary Material for:**

**Safeguarding the chalk to protect salmon genetic diversity…**

**Index**

**Page 2** – Supplementary Table 1 - Details of sampled rivers

**Page 5** – Supplementary Table 2 - Details of SNPs removed from hierarchical chalk and non-chalk datasets

**Page 6** – Supplementary Table 3 - Details of SNP loci found to be under selection

**Page 7** – Supplementary Figure 1 - Results of Bayescan and fdist for determining loci under selection for the full, non-chalk only and chalk only datasets

**Page 8** – Supplementary Figure 2 – Fst outlier loci minor allele frequencies

**Page 9** – Supplementary Figure 3 – Results of hierarchical fdist F_ST_ model for determining loci under selection

**Page 10** – Supplementary Figure 4 – Results of conditioned Redundancy Analysis

**Page 11** – Supplementary Figure 4 - Pairwise F_ST_ between sampled rivers

**Page 12** – Supplementary Figure 5 - Evanno *et al*. (2005) delta *K* (Δ*K* ) results for the hierarchical STRUCTURE analyses of genetic structuring

**Page 13** – Supplementary Figure 6 - Discriminant Analysis of Principal Components (DAPC) analyses for chalk and non-chalk rivers

**Page 14** – Supplementary Figure 7 - Discriminant Analysis of Principal Components (DAPC) analyses for all rivers highlighting two potential strays in French chalk rivers

**Page 15** – Supplementary Figure 8 – Neighbour-joining dendrograms for all, neutral and outlier datasets

**Page 16** – Supplementary Figure 9 - Contribution of rivers and regional groups of rivers to allelic and gene diversity measures

**Supplementary Table 1.** Basic measures of genetic diversity for Atlantic salmon from 42 UK, Irish and French rivers screened for variation at 93 single nucleotide polymorphism (SNP) markers. Risk categories are taken from the NASCO Rivers Database (https://nasco.int/rivers-database/).

| Code | River | Country | River type | N_1_ | Failed samples | Removed full sibs | N_2_ | H_O_ | H_S_ | G_IS_ | G_IS__p | P_POLY SS_ | NASCO risk category |
| --- | --- | --- | --- | --- | --- | --- | --- | --- | --- | --- | --- | --- | --- |
| TYN | Tyne | England | non-chalk | 23 | - | - | 23 | 0.299 | 0.298 | -0.002 | 0.454 | 88.32% | Low |
| TWD | Tweed | England | non-chalk | 24 | 1 | - | 23 | 0.293 | 0.298 | 0.017 | 0.232 | 85.62% | Low |
| WEA | Wear | England | non-chalk | 22 | - | - | 22 | 0.312 | 0.299 | -0.044 | 0.035 | 88.09% | Moderate |
| EDN | Eden | England | non-chalk | 24 | - | 2 | 22 | 0.291 | 0.282 | -0.033 | 0.107 | 86.52% | High |
| LUN | Lune | England | non-chalk | 24 | - | - | 24 | 0.316 | 0.309 | -0.023 | 0.178 | 90.18% | Moderate |
| DEE | Dee | Wales & England | non-chalk | 32 | 3 | - | 29 | 0.312 | 0.320 | 0.027 | 0.109 | 91.19% | High |
| BOY | Boyne | Ireland | non-chalk | 23 | - | 1 | 22 | 0.315 | 0.311 | -0.013 | 0.297 | 92.74% | High |
| AVO | Avoca | Ireland | non-chalk | 46 | - | - | 46 | 0.311 | 0.316 | 0.014 | 0.177 | 91.25% | High |
| ECL | Eastern Cleddau | Wales | non-chalk | 23 | - | 3 | 20 | 0.308 | 0.307 | -0.003 | 0.455 | 87.90% | High |
| TWY | Twyi | Wales | non-chalk | 26 | 4 | 4 | 18 | 0.317 | 0.322 | 0.013 | 0.347 | 89.65% | High |
| USK | Usk | Wales | non-chalk | 47 | 1 | 1 | 45 | 0.309 | 0.316 | 0.022 | 0.119 | 88.99% | Moderate |
| WYE | Wye | Wales & England | non-chalk | 65 | - | - | 65 | 0.312 | 0.317 | 0.014 | 0.164 | 90.18% | High |
| SEV | Severn | Wales & England | non-chalk | 32 | 1 | 1 | 30 | 0.319 | 0.316 | -0.010 | 0.310 | 88.41% | High |
| ELY | East Lyn | England | non-chalk | 25 | 5 | 1 | 19 | 0.277 | 0.274 | -0.013 | 0.309 | 84.70% | High |
| TOR | Torridge | England | non-chalk | 55 | 7 | 4 | 44 | 0.321 | 0.316 | -0.017 | 0.199 | 90.83% | High |
| CAM | Camel | England | non-chalk | 34 | 5 | 1 | 28 | 0.310 | 0.304 | -0.020 | 0.207 | 89.84% | High |
| FOW | Fowey | England | non-chalk | 67 | 1 | 14 | 52 | 0.325 | 0.317 | -0.027 | 0.052 | 90.77% | High |
| LYN | Lynher | England | non-chalk | 35 | 1 | 2 | 32 | 0.321 | 0.313 | -0.026 | 0.106 | 90.19% | Moderate |
| TAM | Tamar | England | non-chalk | 86 | 1 | - | 85 | 0.325 | 0.318 | -0.022 | 0.030 | 91.00% | High |
| TAV | Tavy | England | non-chalk | 48 | - | - | 48 | 0.325 | 0.322 | -0.012 | 0.256 | 91.78% | High |
| PLY | Plym | England | non-chalk | 17 | - | 1 | 16 | 0.310 | 0.300 | -0.032 | 0.145 | 84.95% | High |
| DAV | Devon Avon | England | non-chalk | 23 | - | - | 23 | 0.317 | 0.319 | 0.007 | 0.407 | 93.01% | High |
| DAR | Dart | England | non-chalk | 48 | 1 | 3 | 44 | 0.322 | 0.306 | -0.053 | 0.004 | 88.49% | High |
| TEI | Teign | England | non-chalk | 43 | - | 2 | 41 | 0.312 | 0.312 | 0.000 | 0.526 | 92.32% | Moderate |
| EXE | Exe | England | non-chalk | 49 | - | 1 | 48 | 0.320 | 0.318 | -0.008 | 0.328 | 91.08% | High |
| SIE | Sienne | France | non-chalk | 40 | 1 | - | 39 | 0.297 | 0.313 | 0.050 | 0.005 | 90.89% | Moderate |
| SEE | Sée | France | non-chalk | 40 | - | - | 40 | 0.306 | 0.310 | 0.013 | 0.226 | 90.32% | Moderate |
| TRI | Trieux | France | non-chalk | 40 | - | - | 40 | 0.304 | 0.310 | 0.021 | 0.136 | 89.96% | Moderate |
| LEG | Léguer | France | non-chalk | 40 | - | - | 40 | 0.303 | 0.303 | -0.001 | 0.477 | 88.20% | Low |
| ELN | Elorn | France | non-chalk | 40 | 2 | - | 38 | 0.300 | 0.298 | -0.007 | 0.370 | 87.59% | Moderate |
| SCO | Scorff | France | non-chalk | 36 | - | - | 36 | 0.284 | 0.298 | 0.047 | 0.020 | 87.61% | Low |
| ADR | Adour | France | non-chalk | 22 | - | - | 22 | 0.290 | 0.300 | 0.035 | 0.077 | 83.60% | Moderate |
| FRO | Frome | England | chalk | 39 | 1 | 1 | 37 | 0.297 | 0.299 | 0.008 | 0.369 | 84.29% | High |
| PID | Piddle | England | chalk | 36 | - | - | 36 | 0.281 | 0.289 | 0.030 | 0.059 | 86.88% | High |
| STO | Dorset Stour | England | chalk | 87 | 4 | 8 | 75 | 0.299 | 0.287 | -0.040 | 0.003 | 81.69% | High |
| HAV | Hampshire Avon | England | chalk | 36 | - | 2 | 34 | 0.286 | 0.286 | 0.000 | 0.468 | 84.68% | High |
| TES | Test | England | chalk | 43 | - | - | 43 | 0.292 | 0.288 | -0.012 | 0.267 | 86.22% | High |
| ITC | Itchen | England | chalk | 45 | - | - | 45 | 0.290 | 0.297 | 0.022 | 0.124 | 87.15% | High |
| MEO | Meon | England | chalk | 53 | - | 22 | 31 | 0.276 | 0.291 | 0.052 | 0.007 | 84.54% | Not Determined |
| ARQ | Arques | France | chalk | 40 | 1 | - | 39 | 0.329 | 0.338 | 0.026 | 0.088 | 93.06% | High |
| BRE | Bresle | France | chalk | 64 | 2 | 1 | 61 | 0.331 | 0.330 | -0.002 | 0.450 | 92.24% | Moderate |
| CAN | Canche | France | chalk | 40 | - | - | 40 | 0.351 | 0.353 | 0.004 | 0.405 | 93.43% | High |
| Total |  |  |  | 1682 | 42 | 75 | 1565 |  |  |  |  |  |  |

N_1_ – number of individuals successfully genotyped; N_2_ – number of individuals after removal of failed samples and full sibs (Yank-2); A_R_ – allelic richness, with 95% confidence intervals in parentheses; H_O_ – observed heterozygosity; H_S_ – expected heterozygosity; G_IS_ – inbreeding coefficient; G_IS__p – p value for G_IS_; P_POLY SS_ – percentage of polymorphic loci after subsampling to the smallest sample size.

**Supplementary Table 2.** Details of SNP loci removed from the non-chalk and chalk hierarchical datasets. River codes are as given in Table 1.

| **Dataset** | **Locus** | **Reason** |
| --- | --- | --- |
| Non-chalk only | Ssa_58789 | Monomorphic in non-chalk rivers except for single heterozygote in DAV |
|  | Ssa_24091 | Monomorphic in non-chalk rivers except for three heterozygotes (2x SEE, 1x SIE) |
|  | Ssa_18895 | Monomorphic in non-chalk rivers except for nine heterozygotes (2x WYE, 1x USK, 1x SEV, 5x BOY) |
|  | Ssa_69001 | Monomorphic in non-chalk rivers except for two heterozygotes in BOY |
| Chalk only | Ssa_28584 | Monomorphic in all chalk rivers |
|  | Ssa_73710 | Monomorphic in chalk rivers except for three heterozygotes (1x PID, 1x STO, 1x CAN) |
|  | Ssa_21426 | Monomorphic in chalk rivers except for ten heterozygotes (2x ITC, 2x ARQ, 3x BRE, 3x CAN) |
|  | Ssa_72732 | Monomorphic in chalk rivers except for four heterozygotes (1x FRO, 1x STO, 1x TES, 1x MEO) |
|  | Ssa_13920 | Monomorphic in chalk rivers except for eight heterozygotes (5x ITC, 3x MEO) |

**Supplementary Table 3.** Details of genome location for loci found to be under selection in F_ST_-based outlier tests (Bayescan, fdist and hierarchical fdist) and associated with environmental variables in Redundancy Analyses (RDA).

| **Locus** | **LG^a^** | **SNP position^b^** | **Type** | **Gene** | **Gene abbreviation** | **Test^c^** |
| --- | --- | --- | --- | --- | --- | --- |
| Ssa_76064 | ssa01 | 46176517 | intron | neurexin 3a | nrxn3a | fdist |
| Ssa_30724 | ssa01 | 85159308 | non-coding |  |  | RDA2 |
| Ssa_57180 | ssa05 | 39399608 | intron | transmembrane and coiled-coil domains 6 | tmco6 | fdist |
| Ssa_six6 | ssa09 | 23820071 | non-coding |  |  | Bayescan |
| Ssa_62568 | ssa11 | 87826218 | psuedogene |  | LOC106563332 | fdist, hierarchical fdist |
| Ssa_24091 | ssa13 | 47801856 | intron | serine/threonine-protein kinase 35 | stk35 | fdist |
| Ssa_69865 | ssa13 | 78813823 | non-coding |  |  | Bayescan, fdist, hierarchical fdist |
| Ssa_67740 | ssa14 | 41659245 | intron | muscleblind-like protein 1 | mbnl1 | RDA1 & RDA2 |
| Ssa_58789 | ssa15 | 39050401 | intron | insulin-like growth factor 2 receptor | igf2r | fdist |
| Ssa_87179 | ssa16 | 4880999 | intron | uncharacterised gene | LOC106573079 | RDA2 |
| Ssa_73197 | ssa18 | 6835282 | intron | Kv channel-interacting protein 2-like | kchip2 | fdist |
| Ssa_55742 | ssa21 | 38896843 | non-coding |  |  | fdist |
| Ssa_18895 | ssa24 | 31184582 | intron | disabled homolog 2-interacting protein-like | dab2ipa | fdist |
| Ssa_55100 | ssa24 | 29841615 | intron | neurotrophic tyrosine kinase receptor, type 2a | ntrk2a | fdist |
| Ssa_34412 | ssa29 | 34045134 | intron | vav guanine nucleotide exchange factor 3 | vav3 | fdist |

^a^ LG – Linkage Group; ^b^ position of SNP based on alignment to Ssal_v3.1 Atlantic salmon genome; ^c^ RDA1 – unconditioned RDA; RDA2 – RDA conditioned by geographic distance from most northernly river

**Supplementary Figure 1.** Results for selection tests for chalk-only and non-chalk-only datasets. a) - b) Bayescan results. The dashed blue line represents the log_10_ PO value above which selection on loci is considered significant. c) – d) fdist results. Dashed lines are the upper and lower 99% confidence level of the simulated neutral distribution. Loci under either divergent or balancing selection are shown in red with loci under divergent selection labelled with locus name. a) & c) results for the non-chalk river data set of 89 SNPs screened in 32 rivers; b) & d) results for the chalk river data set of 88 SNPs screened in 10 rivers.

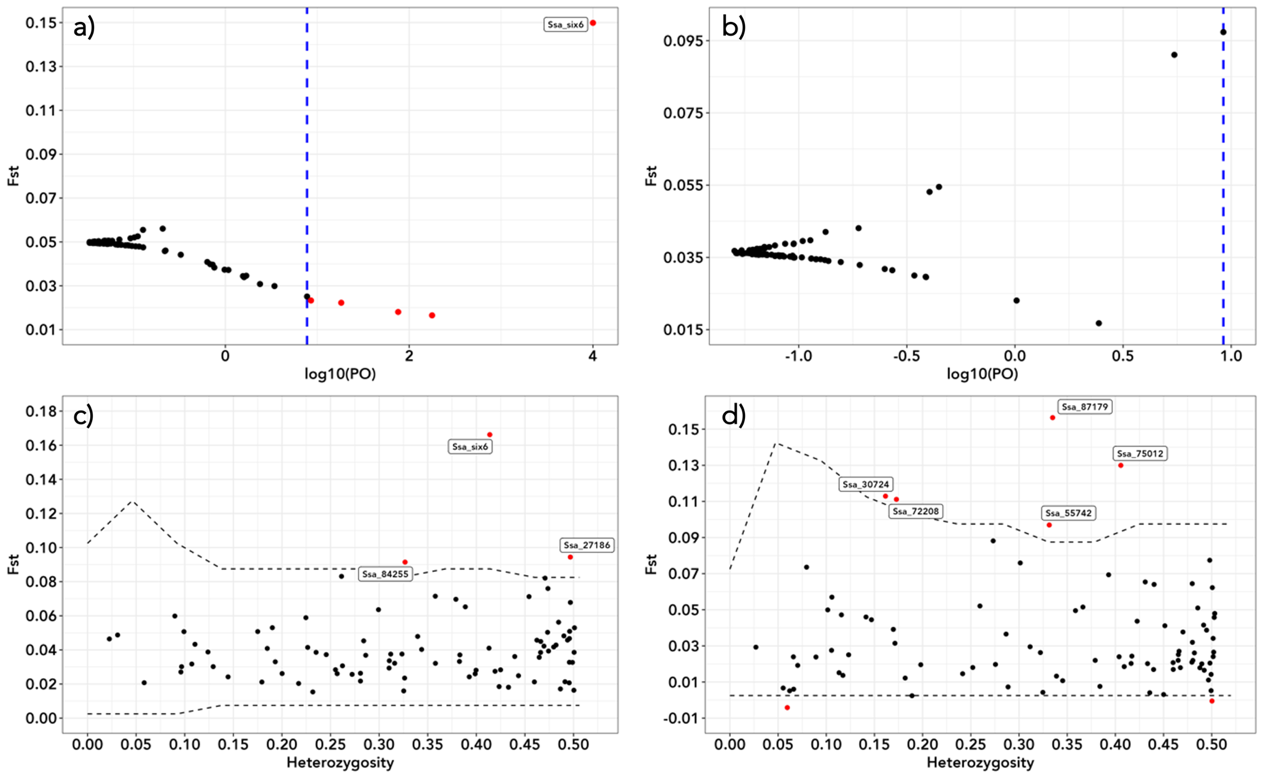

**Supplementary Figure 2** Heatmap showing the minor allele frequencies for 15 SNPs found to be under divergent selection in outlier tests and associated with environment in Redundancy Analyses (RDA). Numbers above each column denotes in which test each locus was identified (1 – Bayescan, 2 – fdist, 3 – hierarchical fdist, 4 – unconditioned RDA, 5 – RDA conditioned by geographical distance from northern-most river)

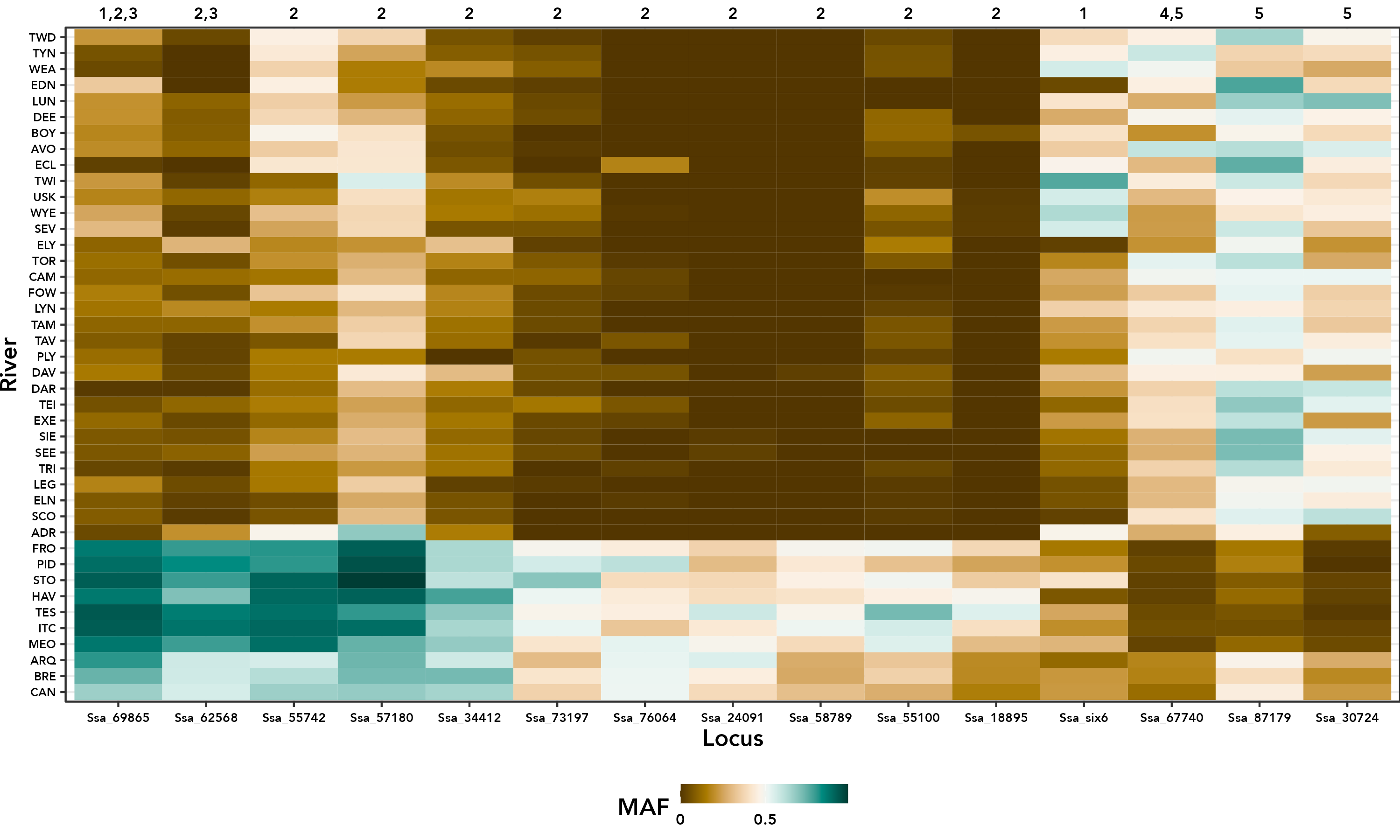

**Supplementary Figure 3.** Results of hierarchical fdist F_ST_ model based on two groupings of rivers. a) chalk v non-chalk and b) UK chalk v FR chalk v UK non-chalk v FR non-chalk. Dashed lines are the upper and lower 99% confidence level of the simulated neutral distribution.

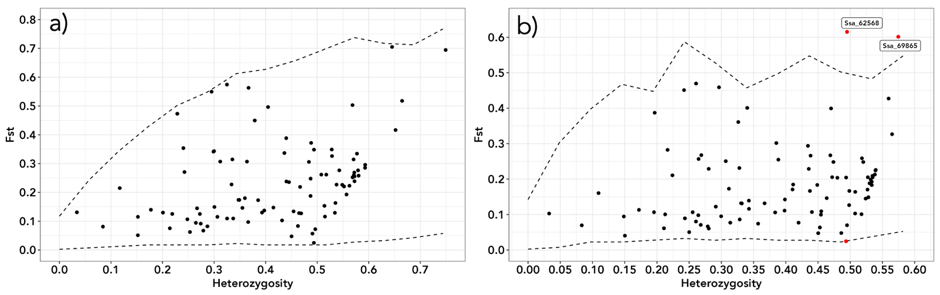

**Supplementary Figure 4.** Results of Redundancy Analysis conditioned on geographic distance from northern-most river. a) Plot of conditioned redundancy analysis axes 1 v 2. Black vectors represent loadings for the seven retained bioclim variables. River codes are as given in Supplementary Table 1. b-d) Plot of minor allele frequency for loci Ssa_67740 & Ssa_30724 versus standardised bio17 (precipitation of the driest quarter) values and locus Ssa_87179 versus standardised bio10 (mean temperature of the warmest quarter) values. The red line represents the linear regression, and the grey shaded area is the 95% confidence interval.

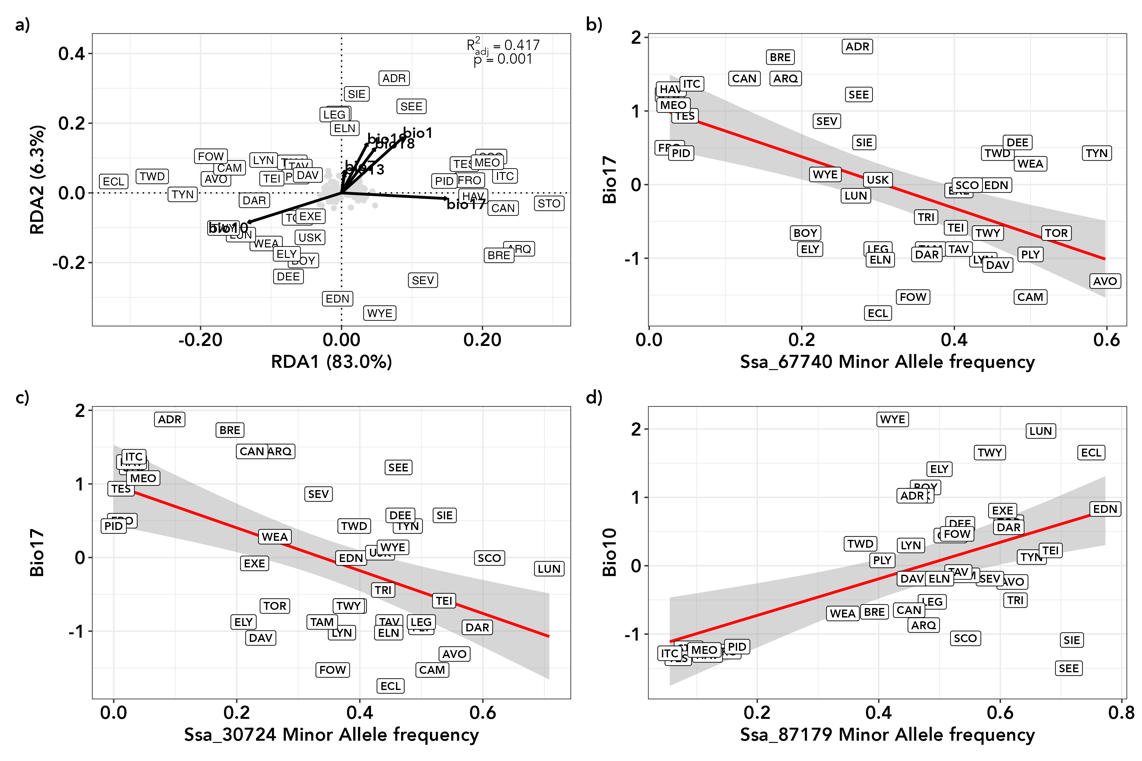

**Supplementary Figure 5.** Heatmap showing pairwise Weir & Cockerman’s F_ST_ between 42 British, Irish and French Atlantic salmon rivers. River codes are as given in Supplementary Table 1.

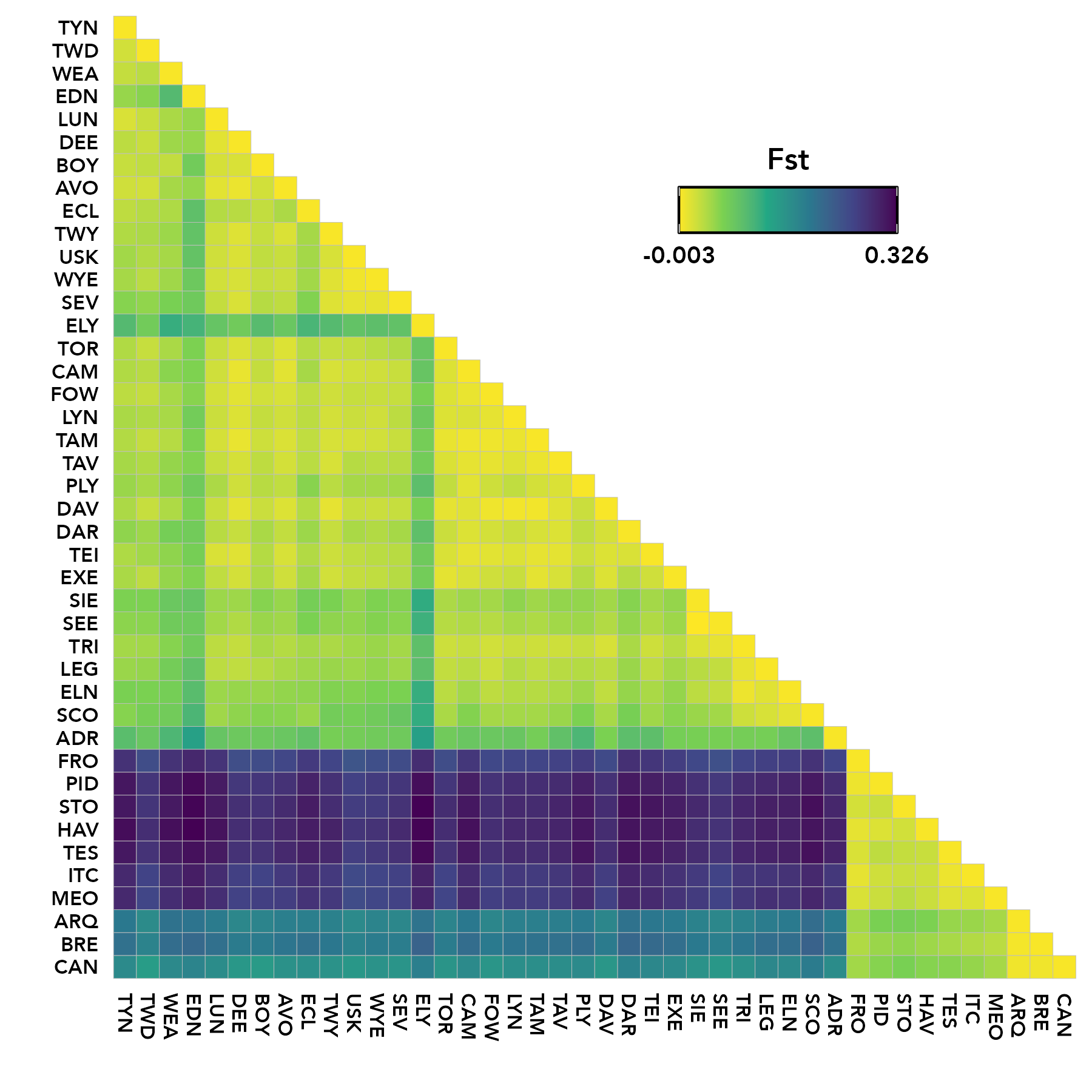

**Supplementary Figure 6.** Results for Evanno *et al*. (2005) delta *K* (Δ*K* ) for the hierarchical STRUCTURE analyses of genetic structuring in 42 Atlantic salmon rivers.

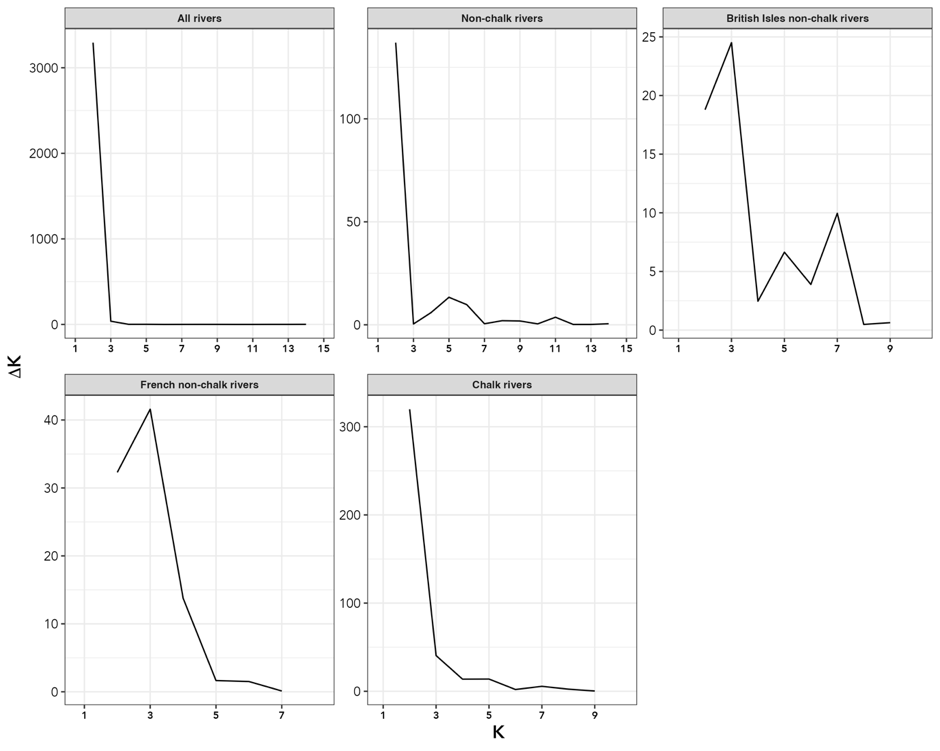

**Supplementary Figure 7.** Discriminant Analysis of Principal Components (DAPC) analyses for Atlantic salmon from UK, Irish and French rivers. Each dot represents a sampled individual fish. River codes are as given in Supplementary Table 1 and dots are coloured as given in Figure 2. a) – non-chalk river data set (1124 individuals from 32 rivers) and b) - chalk river data set (441 individuals from 10 rivers).

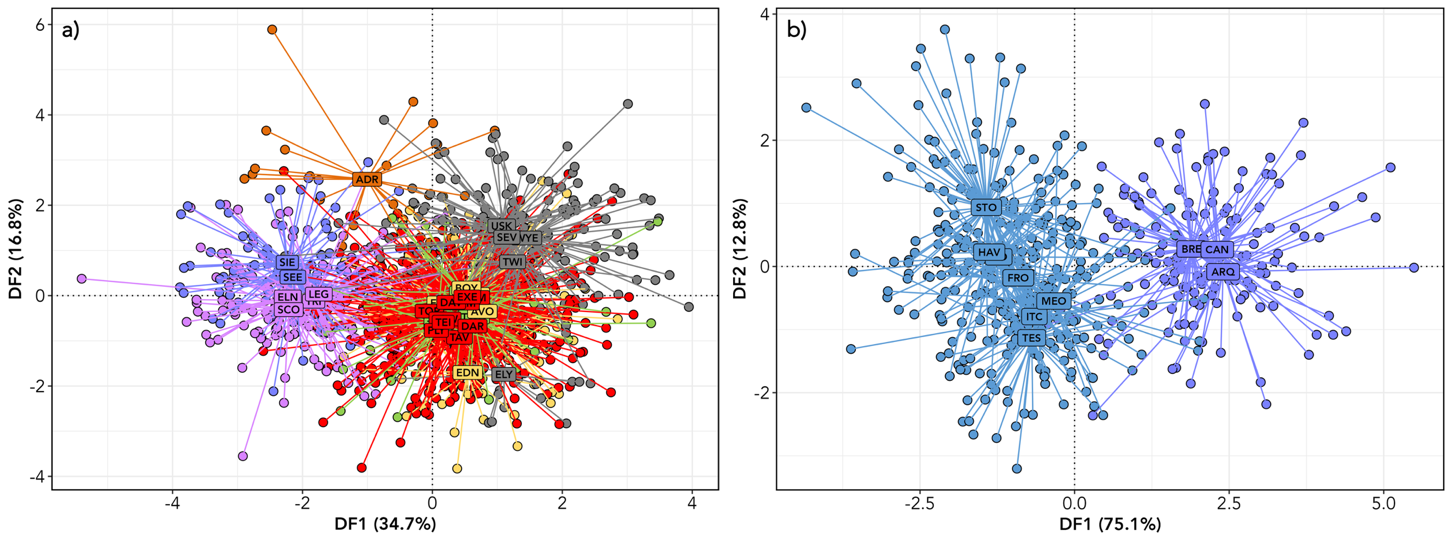

**Supplementary Figure 8.** Discriminant Analysis of Principal Components (DAPC) analyses for Atlantic salmon from 42 UK, Irish and French rivers highlighting the two suspected strays (CAN30 & ARQ11) from non-chalk rivers into two French chalk rivers. Each dot represents a sampled individual fish. River codes are as given in Supplementary Table 1 and dots are coloured as given in Figure 2.

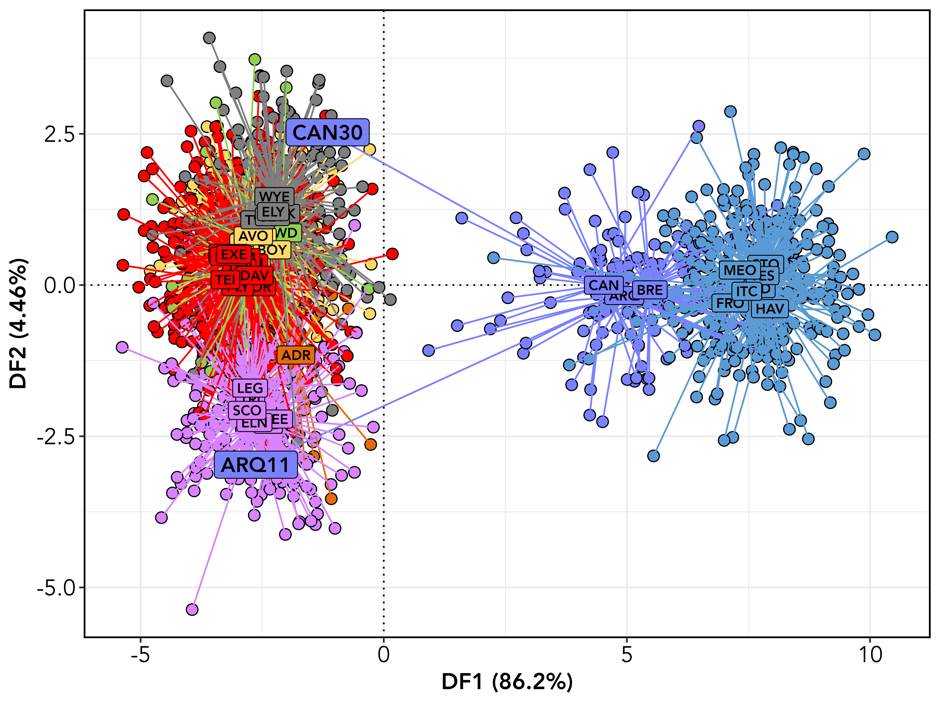

**Supplementary Figure 9.** Population-based Neighbour-joining dendrograms for a) all, b) neutral and c) outlier datasets. River codes are as given in Supplementary Table 1 and branches are coloured as given in Figure 2.

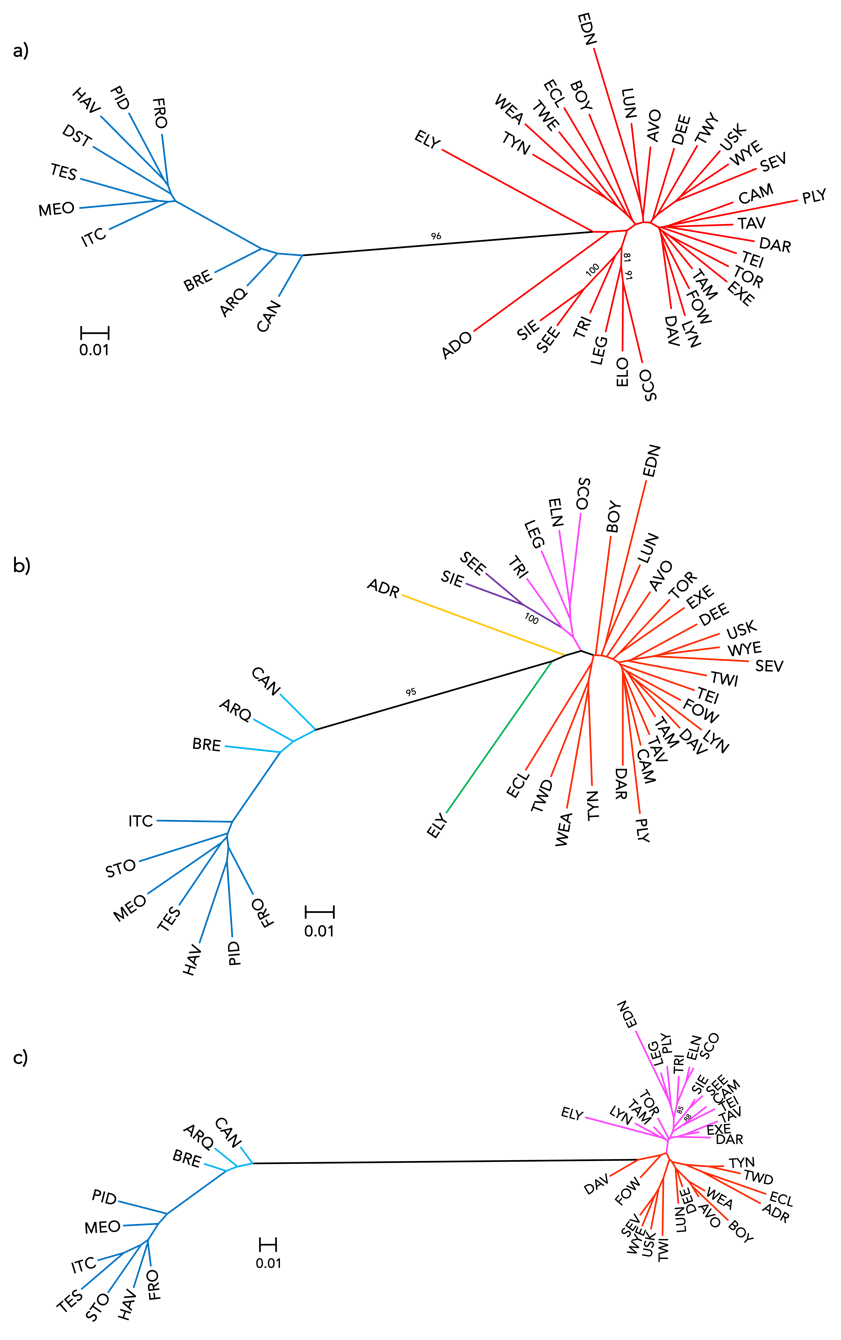

**Supplementary Figure 10.** Contribution of 42 Atlantic salmon rivers and four major regional groups of rivers to genomic diversity. Contributions are partitioned into within-population/group (*A_S_* and *H_S_*), between-population/group (*D_A_* and *D_G_*) and total (*A_T_* and *H_T_*) components of allelic and gene diversity, respectively.

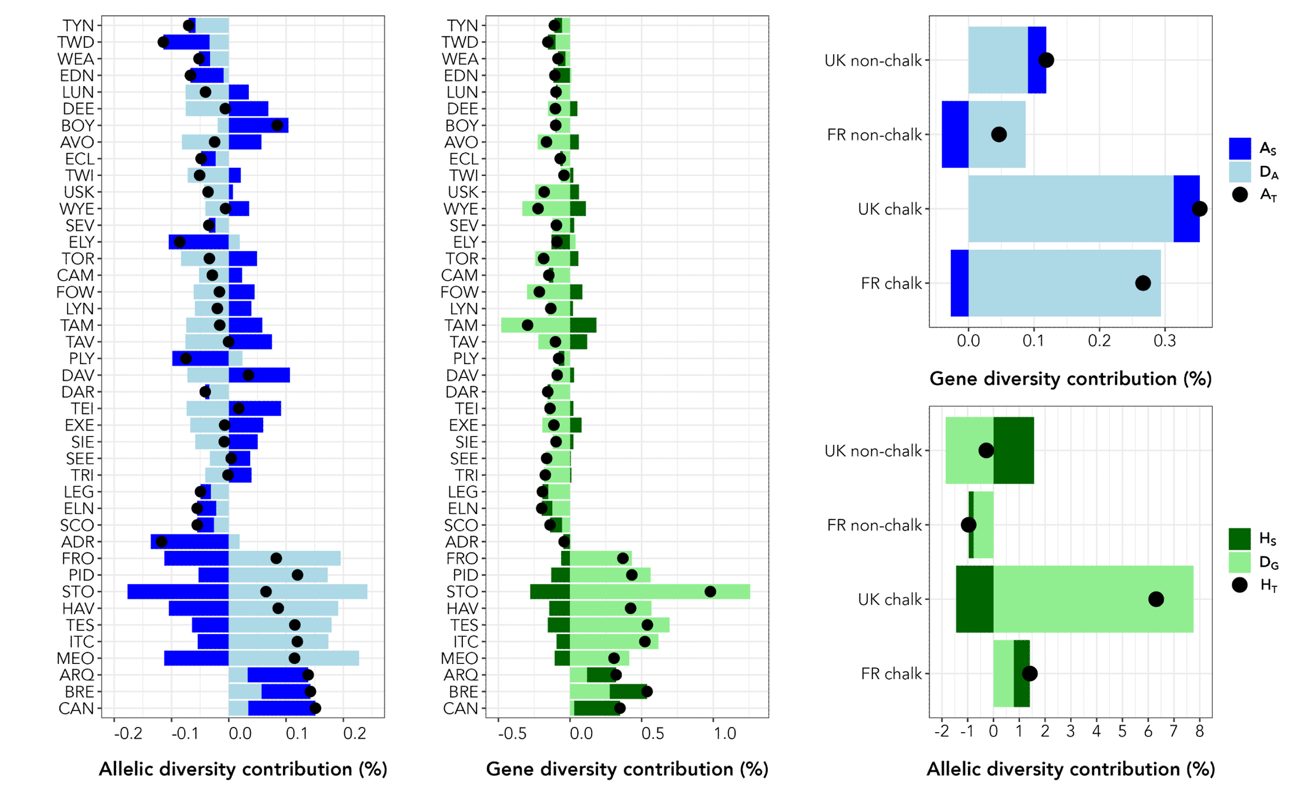
